# Rapid radiation of a plant lineage sheds light on the assembly of dry valley biomes

**DOI:** 10.1101/2024.05.05.592625

**Authors:** Ya-Ping Chen, Purayidathkandy Sunojkumar, Robert A. Spicer, Richard G.J. Hodel, Douglas E. Soltis, Pamela S. Soltis, Alan J. Paton, Miao Sun, Bryan T. Drew, Chun-Lei Xiang

**Affiliations:** CAS Key Laboratory for Plant Diversity and Biogeography of East Asia, Kunming Institute of Botany, Chinese Academy of Sciences, Kunming, 650201, China; Department of Botany, University of Calicut, Kerala, 673635, India; School of Environment, Earth and Ecosystem Sciences, The Open University, Milton Keynes, MK7 6AA, UK; CAS Key Laboratory of Tropical Forest Ecology, Xishuangbanna Tropical Botanical Garden, Chinese Academy of Sciences, Mengla, 666303, China; State Key Laboratory of Tibetan Plateau Earth System, Resources and Environment (TPESRE), Institute of Tibetan Plateau Research, Chinese Academy of Sciences, Beijing, 100101, China; Department of Botany, National Museum of Natural History, MRC 166, Smithsonian Institution, Washington, DC 20013, USA; Data Science Lab, Office of the Chief Information Officer, Smithsonian Institution, Washington, DC 20560, USA; Florida Museum of Natural History, University of Florida, Gainesville, FL 32611, USA; Department of Biology, University of Florida, Gainesville, FL 32611, USA; Royal Botanic Gardens, Kew, Richmond, TW9 3AE, UK; National Key Laboratory for Germplasm Innovation & Utilization of Horticultural Crops, Huazhong Agricultural University, Wuhan, 430070, China; Department of Biology, University of Nebraska-Kearney, Kearney, NE 68849, USA

**Keywords:** drought tolerance, East Asian monsoon, Hengduan Mountains, Tibetan Plateau, woodines

## Abstract

Southwest China is characterized by high plateaus, large mountain systems, and deeply incised dry valleys formed by major rivers and their tributaries. Despite the considerable attention given to alpine plant radiations in this region, the timing and mode of the diversification of the numerous plant lineages in the dry valley habitat remains unknown. To address this knowledge gap, we investigate the macroevolution of *Isodon* (Lamiaceae), a lineage commonly distributed in the dry valleys in southwest China and wetter areas of Asia and Africa. We reconstructed a robust phylogeny encompassing nearly 90% of the approximately 140 extant *Isodon* species using transcriptome and genome-resequencing data. Our results suggest a rapid radiation of *Isodon* during the Pliocene that coincided with a habit shift from herbs to shrubs and a habitat shift from humid areas to dry valleys. The shrubby growth form likely acted as a preadaptation allowing for the movement of *Isodon* species into these valleys. Ecological analysis highlighted aridity and precipitation as key factors influencing the niche preferences of different growth forms and species richness of *Isodon*. Integrating our results with insights from tectonic movements in the Tibetan Plateau and adjacent regions, we infer that the interplay between topography and the evolution of the East Asian monsoon since the middle Miocene likely contributed to the formation of the dry valley biome in southwest China. This study enhances our understanding of evolutionary dynamics and ecological drivers shaping the distinctive flora of this region.

## Introduction

Montane plant radiations have attracted considerable interest, especially for the alpine floras of the Andes and Hengduan Mountains (1−4). However, compared to the evolutionary trajectories of high elevation clades, little is known about the diversification of lineages in the adjacent inter-montane valleys. Southwest China is characterized by high plateaus (Tibetan Plateau, Yunnan Plateau), large mountain systems (Himalaya, Hengduan Mountains), and deeply incised valleys formed by the Yarlung Zangbo River (YZR, the upper reaches of the Brahmaputra River), Nujiang (the upper reaches of the Salween River), Lancangjiang (the upper reaches of the Mekong River), Jinshajiang (the upper reaches of the Yangtze River), and Yuanjiang (the upper reaches of the Red River) and their tributaries. In the Hengduan Mountains, many of the rivers flow southward in parallel and the intense rain shadows cast by the high mountains have resulted in valleys dominated by semi-arid or arid climates with savanna-like (“savanna of valley type”) or maquis-like (“maquis of valley type”) vegetation (5, 6). Over 3,000 seed plant species have been documented from the dry valleys in southwest China, with nearly one-third of them endemic to this region (5−7). As a unique biome, this river valley ecosystem has been shown to be at risk due to climate change and increasing intensity of human activities (8, 9). The above rivers may have once been tributaries of a single, southward flowing river system—the paleo-Red River (PRR), which drained into the South China Sea (10, 11). Major headwater river capture events from the huge PRR might have happened later, forming the modern drainage system pattern (10, 11).

These events were likely driven by tectonic movements of the Tibetan Plateau, Himalaya, and Hengduan Mountains (collectively referred to as the THH region) and/or the intensification of the East Asian monsoon. Whether the river capture events have been involved in the development of modern drainages and the timing of disruption of the old drainage system is still a matter of considerable debate (10−14). The effects of drainage reorganization on phytogeographic patterns (summarized in ref. 15) and the genetic diversity within endangered species endemic to dry valleys (summarized in ref. 8) have been thoroughly investigated, but it is still unclear when the dry valley plants originated and how they diversified in this specific habitat.

Dryland biomes, which span over 40% of Earth’s land surface, are one of the most threatened and vulnerable ecosystems, despite being overlooked in biodiversity research (16−19). An accelerated dryland expansion and degradation is anticipated as a consequence of climate change and the burgeoning human population (20, 21). Understanding the evolutionary dynamics underlying the development of drought-avoidance functional traits and their correlation with arid conditions is essential for informing conservation practices and devising effective adaptation strategies.

Previous studies have revealed various strategies employed by plants and animals to shift into environments analogous to dry valleys in other regions, such as the tropical seasonally dry forests and savannas (e.g., ref. 22−24). However, little is known about the traits that facilitate adaptation to dry habitats in mountain systems (but see ref. 25). Indeed, diversification prompted by emerging ecological opportunities resulting from climate change or mountain building frequently necessitates adaptation across a suite of clade-specific life-history traits (23). The evolutionary transition from herbaceousness to woodiness on islands, i.e., insular woodiness, represents one of the most iconic features of island plant radiations (26−29). Similar accelerated rates of growth-form evolution have also emerged as a common feature of tropical alpine radiations where secondary woodiness or perenniality (shifts from annual to perennial habit) served as a key innovation, enabling lineages to take advantage of ecological opportunities of new cooler, high-elevation habitats provided by the uplift of mountains and facilitating rapid species diversification and evolution of growth forms (1, 30−33). However, these striking parallels between island and island-like alpine radiations have rarely been reported from the floras of the Himalaya and Hengduan Mountains (but see ref. 34). An increasing number of studies employing phylogenomic data have shed light on the factors driving rapid radiations in southwest China (e.g., ref. 35−37), but our understanding of the detailed trajectories of diversification and potential rapid evolutionary radiations, particularly in dry valleys, is still hindered by lack of robust and well-sampled phylogenies and detailed diversification analyses (1).

*Isodon* (Schrad. ex Benth.) Spach (Ocimeae, Nepetoideae, Lamiaceae) is a genus of 100–150 species mainly distributed in subtropical to tropical Asia, with two disjunct species endemic to Africa (38–40). The genus is most diverse in southwest China, particularly in the dry valleys in the Hengduan Mountains, which is considered the center of diversity for the genus (41–43). Species of *Isodon* can be found in nearly all major river valleys of southwest China, making it an ideal lineage to explore the origin and diversification of plants adapted to these dry valley systems.

Notably, *Isodon* species showing contrasting growth forms (shrub vs. herb) are usually distributed in distinct habitats (dry vs. wet). Previous biogeographic investigations failed to clarify the spatiotemporal patterns of evolution of the genus due to limited phylogenetic resolution (42). In this study, we built a well-resolved and time-calibrated phylogenomic framework for the genus using transcriptome and whole-genome resequencing data. We further conducted a series of biogeographic, macroevolutionary, and macroecological analyses to explore the trajectories of plant diversification in dry valleys in southwest China and test whether a shrubby habit facilitated colonization and speciation in these dry valley systems spatially and temporally, using *Isodon* as an example. In a macroevolutionary context, we hypothesize that (H1) *Isodon* experienced a recent rapid radiation, and the diversification rate shift is correlated with a transition in growth form from herbs to shrubs and in habitat from humid area to dry valleys. We also hypothesize that (H2) shrubby growth form and/or dry valley habitat triggers elevated speciation rates in *Isodon*. From a macroecological perspective, we hypothesize that (H3) environmental factors related to increasing aridity and lower precipitation are major drivers differentiating the distribution of herbs and shrubs and impacting the species richness of *Isodon*. Finally, combining all evidence, we explore the formation of dry valley biomes in southwest China, and its implications for our understanding of dryland biomes globally.

## Results

### Phylogenetic Relationships within *Isodon*

We sampled a total of 171 accessions of 126 taxa from all major regions across the distribution of *Isodon* in Asia and Africa, and from major river valleys in southwest China (*SI Appendix*, Figs. S1 and S2), covering ca. 90% of extant *Isodon* species. An average of 95,320 transcripts were de novo assembled for the 149 transcriptomes with an average N50 scaffold length of 2,100 base pairs (bp). After removing 133 low-complexity sequences and 195 sequences with at least one unexpected stop codon, we used the remaining 35,108 coding sequences (CDS) filtered from the genome of *I. rubescens* (Hemsl.) H. Hara as the target file for assembly of the 30 genome resequencing samples. The number of orthologs retrieved for each of the 180 samples ranged from 3,936 (*I. phyllopodus* (Diels) Kudô b) to 6,118 (*Siphocranion macranthum* (Hook. f.) C.Y. Wu) with an average of 5,135. For the 7,204 orthologs used in the phylogenetic analyses, the aligned lengths ranged from 300 bp to 10,917 bp with an average of 1,171 bp. The concatenated dataset consisted of 8,437,966 columns with an overall matrix occupancy of 62%. A summary of the assembly statistics is shown in Data S1.

Previous phylogenetic studies using several molecular markers and plastome sequences consistently recovered four well-supported clades (Clade I−Clade IV) within *Isodon*, but the phylogenetic relationships within Clade IV, which comprises ca. 80% species of the genus, were poorly resolved (42, 44). Compared with previous analyses, both the ML tree and the coalescent tree retrieved in present study had greater resolution, with most nodes receiving maximal support (Fig. 1, *SI Appendix*, Figs. S3 and S4: ultrafast bootstrap (UFBS) = 100%, local posterior probability (LPP) = 1.00). Our study also supported the monophyly of the four clades detected previously. Clade II, which is comprised of the two species endemic to Africa, was sister to Clade I, and clade I-II was sister to clade III-IV. Clade III comprised the three accessions of *I. ternifolius* (D. Don) Kudô. A large majority of *Isodon* were grouped in Clade IV, within which we further delimited four subclades (Clade IVa−Clade IVd).

**Fig. 1.**
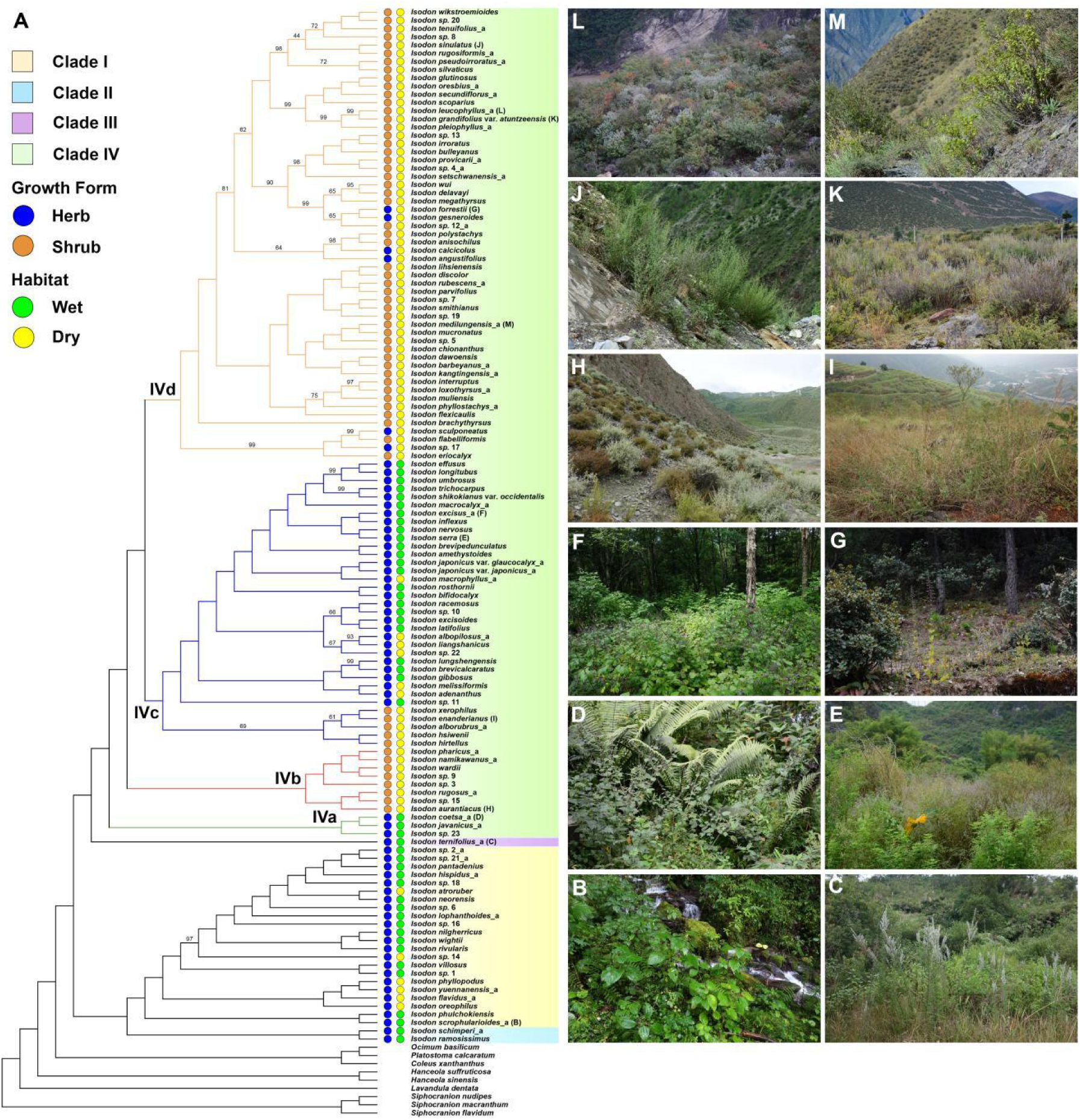
Phylogeny and growth-form and habitat diversity of *Isodon*. (A) Cladogram of the maximum-likelihood tree of *Isodon* estimated using the concatenation method of IQ-TREE based on the dataset of 7,204 low-copy nuclear genes. Multiple accessions of species are removed. Numbers on nodes indicate ultrafast bootstrap (UFBS) values < 100%. The letters in brackets following species names correspond to the images on the right (B–M). (B) *Isodon scrophularioides* from the southern Himalaya. (C) *Isodon ternifolius* from southern Yunnan, China. (D) *Isodon coetsa* from the southern Himalaya. (E) *Isodon serra* from the limestone area of south China. (F) *Isodon excisus* from northeast China. (G) *Isodon forrestii* in the forest of *pinus yunnanensis* in southwest China. (H) *Isodon aurantiacus* in the dry valley of the Yarlung Zangbo River. (I) *Isodon xerophilus* in the dry valley of Yuanjiang. (J) *Isodon sinulatus* in the dry valley of Nujiang. (K) *Isodon grandifolius* var. *atuntzeensis* in the dry valley of Lancangjiang. (L) *Isodon leucophyllus* from the Tiger Leaping Gorge in the dry valley of Jinshajiang. (M) *Isodon medilungensis* in the dry valley of Yalongjiang (a tributary of Jinshajiang).

The topologies of the ML tree and coalescent tree of *Isodon* were largely congruent, except for the relationships within Clade IVd, where most of the conflicts between the two topologies were found (*SI Appendix*, Fig. S5). Many nodes in this clade were poorly resolved with low support values (Fig. 1, *SI Appendix*, Figs. S3, S4: UFBS < 95%, LPP < 0.95) and extremely short branch lengths. The quartet sampling analyses found that quartets were largely informative at all nodes with a mean quartet informativeness (QI) of 0.998 (*SI Appendix*, Fig. S6). The mean quartet concordance (QC) for the branches was 0.64. A total of 12 nodes (6.7%) had QC values below -0.05 (i.e., counter-support for an alternative quartet topology), and all these nodes corresponded to short branches. The polytomy test revealed that seven nodes within *Isodon* were hard polytomies (*p* > 0.05), and all these nodes were located within Clade IV, and especially in Clade IVd (*SI Appendix*, Fig. S7). Most of the above 12 nodes with QC < -0.05 rejected the null hypothesis of a polytomy, but the quartet support calculated by ASTRAL was low at these nodes (e.g., 36% at node A, with alternative quartet support 35% and 29%; *SI Appendix*, Fig. S8). Given the unequal quartet frequencies for alternative topologies at these nodes, the topological conflicts could not only be explained by incomplete lineage sorting, but might also be caused by other factors such as hybridization and/or gene introgression.

For the PhyloNet analyses among major clades of *Isodon*, the optimal inferred phylogenetic network with the highest log pseudo-likelihood included three reticulation events (*SI Appendix*, Fig. S9). Clade II was descended from a putative ancient hybridization event between the common ancestor of Clade I and the common ancestor of clade III-IV, whereas *I. pharicus* (Prain) Murata was of hybrid origin resulting from reticulation between the common ancestor of clade IVc-IVd and *I. rugosus* (Wall. ex Benth.) Codd. Within Clade I, a phylogenetic network with three reticulation events was also optimal with highest log pseudo-likelihood score (*SI Appendix*, Fig. S9). *Isodon scrophularioides* (Wall. ex Benth.) Murata was inferred to be a hybrid between *I. phulchokiensis* (Murata) H. Hara and the common ancestor of the *I. oreophilus* (Diels) A.J. Paton & Ryding ∼ *I. flavidus* (Hand.-Mazz.) H. Hara clade, whereas *I. flavidus* was a hybrid with ancestry contribution from *I. yuennanensis* (Hand.-Mazz.) H. Hara. The third reticulation event was found among the remaining taxa of Clade I.

### Divergence Time and Ancestral Range Estimation

The divergence times estimated using different methods were largely congruent (< 1 million year difference for most nodes) (*SI Appendix*, Figs. S10 and S11). For example, the estimated crown age of *Isodon* was 14.63 million years ago (Ma) (95% highest posterior density (HPD): 10.44−19.35 Ma) in the BEAST chronogram, and 15.01 Ma in the dated treePL phylogeny, both corresponding to the middle Miocene. For brevity, we describe in detail only the results from BEAST in subsequent sections (*SI Appendix*, Fig. S10). Clade II diverged from Clade I at 12.71 Ma (95% HPD: 8.97−16.95 Ma) in the middle to late Miocene, and the crown ages were 5.14 Ma (95% HPD: 3.67−6.80 Ma) and 7.48 Ma (95% HPD: 4.69−10.53 Ma) for Clade I and Clade II, respectively. The split between Clade III and Clade IV was estimated at 8.28 Ma (95% HPD: 5.50−11.51 Ma) in the late Miocene to early Pliocene. Subsequently, all four subclades of Clade IV diverged within 1.4 million years in the Pliocene (from 4.69 Ma to 3.28 Ma) and started to diversify at 3.63 Ma (Clade IVa, 95% HPD: 2.45−4.96 Ma) to 3.09 Ma (Clade IVd, 95% HPD: 2.26−3.97 Ma).

The DEC model was selected as the best model (*SI Appendix*, Table S1) for the BioGeoBEARS analysis. According to the ancestral range estimation (Fig. 2), *Isodon* widely expanded across the Himalaya, Hengduan Mountains, and South and Southeast Asia in the middle Miocene, but it was difficult to determine exactly where the genus originated. It should not have been present in Africa during this time due to the hybrid origin of the African Clade II. Within *Isodon*, a vicariance event separating Clade I and Clade II was later hypothesized, and the ancestral area of Clade I was obscured. Clade IV originated in the Himalaya or the Hengduan Mountains (including the Yunnan-Guizhou Plateau). Subsequently, a vicariance event was hypothesized, leading to the divergence of Clade IVb and clade IVc-IVd. All species of Clade IVb were distributed in the Himalaya, especially in the dry valleys in the YZR. The ancestral area of Clade IVc was inferred to be the Hengduan Mountains, with later dispersal to subtropical China, northern China, the Russian Far East, the Korean Peninsula, and subsequently to Japan. All *Isodon* species endemic to Japan originated from a single vicariance event from subtropical China. The most recent common ancestor (MRCA) of Clade IVd shared the same distribution area as that of Clade IVc, and most species from this clade were endemic to the dry valleys in the Hengduan Mountains. The results of 50 biogeographic stochastic mappings (BSMs) showed that over half of evolutionary events in *Isodon* (65%) involved taxa having undergone speciation within a single area, i.e., narrow sympatry (*SI Appendix*, Table S2). Dispersal (20%) also played an important role in the evolutionary history of *Isodon*, followed by the processes of subset sympatry (8%) and vicariance (7%).

**Fig. 2.**
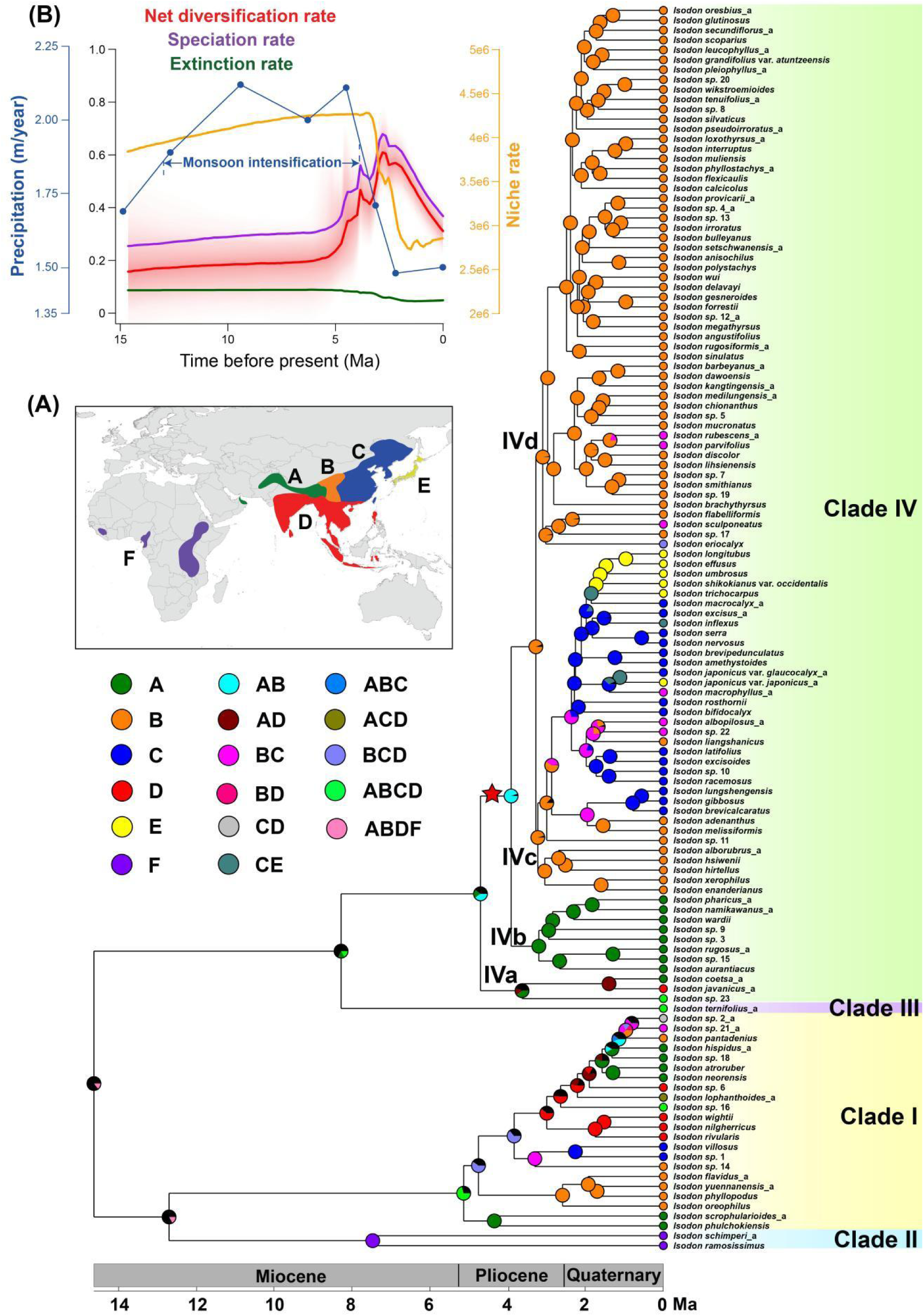
Historical biogeography of *Isodon*. (A) Ancestral range estimation based on the DEC model using BioGeoBEARS. Pie charts at the nodes indicate relative probabilities of all possible geographic ranges. Black color in the circle indicates the sum of probabilities of all alternative ancestral areas with probability <15%. The red star represents major rate shift position on the phylogeny. (B) Net diversification rate (red), speciation rate (blue), extinction rate (green), and evolutionary rates of niche (orange) estimated by BAMM. The unit of net diversification is species/million years, and niche rates are unitless. Monsoon conditions are indicated by the modelled mean annual precipitation for each geologic stage, denoted by blue lines at idealized CO_2_ levels (blue circles) (modified from ref. 83).

### Macroevolutionary Rates

The best shift configuration estimated by Bayesian analysis of macroevolutionary mixtures (BAMM) revealed a rate shift for *Isodon*, occurring at the crown of clade IVb-IVd (*SI Appendix*, Figs. S12 and S13). The average net diversification rate of *Isodon* was estimated as 0.41 species per million years (species/Myr). According to the clade-specific evolutionary rate analyses, the net diversification rates of the clade IVb-IVd was significantly higher than those of the remaining clades, with an average rate of 0.52 (vs. 0.23) species/Myr. The rate-through-time plots suggested that *Isodon* likely underwent a rapid evolutionary radiation during the Pliocene, followed by a decline to lower rates of diversification from the start of the Pleistocene to the present (Fig. 2). The increase in the net diversification rate-through-time curve corresponded to the position of the rate shift in the phylogeny. The CoMET model in TESS also showed similar rate estimates as that of BAMM, with the speciation rate experiencing a rapid acceleration between 5 Ma and 4 Ma and declining sharply from 2 Ma, as well as a relatively constant extinction rate (*SI Appendix*, Fig. S14).

We also explored the association between diversification rates and global paleotemperature and East Asian monsoon by comparing the fit of four diversification models (i.e., constant-rate, time-dependent, temperature-dependent, and monsoon-dependent models) in RPANDA to determine which model better explained the diversification of *Isodon*. The paleoenvironment-dependent analyses found that the exponential speciation without extinction with East Asian monsoon dependence is the best model (*SI Appendix*, Table S3), and the speciation rate of *Isodon* was negatively correlated with East Asian monsoon (*α* = -1.8242). Thus, the speciation rates of *Isodon* gradually increased over time as the intensity of East Asian monsoon declined, and experienced a sharp increase during the Pliocene (*SI Appendix*, Fig. S15), which corroborated the results from BAMM and CoMET.

Rates of ecological niche evolution were generally constant early in the evolution of *Isodon*, but later experienced a strong decrease towards the present, beginning around the late Pliocene (Fig. 2). This shift in the rate of niche evolution corresponded to the crown of Clade IVd (*SI Appendix*, Fig. S16). In addition, the aridity index was the most important variable based on primary loadings of ordinated niche data, followed by variables related to precipitation (*SI Appendix*, Table S4). Notably, the aridity index likely drove the rate shift of niche evolution. When all 39 environmental variables were analyzed together, or when the aridity index was analyzed separately, similar results were observed. However, when the aridity index was excluded, the niche rates gradually decreased through time without any shift (*SI Appendix*, Fig. S17).

### Ancestral State Reconstruction and Correlated Evolution

Ancestral reconstruction of growth form using Mesquite revealed that the ancestor of *Isodon* had a herbaceous growth form (*SI Appendix*, Fig. S18). Following the split of clade I-II and clade III-IV in the middle Miocene, all species of Clade I−Clade III and Clade IVa retained the herbaceous state. Shrubs first evolved from herbs at the MRCA of clade IVb-IVd in the middle Pliocene. All species of Clade IVb and the majority of Clade IVd were shrubs, and the herbaceous growth form was hypothesized to be regained six times within clade IVc-IVd. The most significant shift from shrubs to herbs happened within Clade IVc in the late Pliocene, resulting in a clade of herbaceous species mainly distributed in East Asia.

The evolution of habitat type showed a similar pattern as that of growth form. The ancestral habitat type was estimated as wet area for *Isodon* (*SI Appendix*, Fig. S18). Species adapted to dry habitat first emerged at the MRCA of clade IVb-IVd in the middle Pliocene. Though most species of Clade I were distributed in humid areas, dry valley species evolved three times independently within this clade. Within clade IVb-IVd, lineages with a wet affinity arose only once, all of which were herbs from Clade IVc distributed in East Asia. All species of Clades IVb and IVd and all shrubby species were accustomed to dry valleys.

The Brownian motion (BM) model with the lowest corrected Akaike information criterion (AICc) was selected for niche evolution. As the primary loadings on ordinated niche data were mainly determined by the aridity index, ancestral niche reconstruction (*SI Appendix*, Fig. S19) showed a similar pattern to that of aridity index evolution (*SI Appendix*, Fig. S20). The ancestors of *Isodon* only started to diversify in drier places, possibly the dry valleys, around the crown of clade IVb-IVd.

Our analyses of phylogenetic signal via *D* statistical calculations suggested that both growth form (*D* = -1.577; *D* = 0, *p* = 1.00) and habitat (*D* = -1.321; *D* = 0, *p* = 1.00) displayed a highly conserved pattern of evolution across the *Isodon* phylogeny, indicating that more closely related species were more likely to share these traits. The best model of the correlated evolution tests demonstrated that growth form evolution influenced habitat preference in *Isodon* species but not vice versa (*SI Appendix*, Table S5, Fig. S21). In particular, the transition rates from wet habitat to dry valleys were much higher in shrubby species than in herbaceous ones (*SI Appendix*, Fig. S21).

### Trait-Dependent Analyses

The hidden state speciation and extinction (HiSSE) model analyses found no evidence of a correlation between speciation rate and growth form or habitat states. The best-fit model for both states was the character-independent diversification (CID) model with four hidden states (CID-4) and equal extinction fraction and transition rate (*SI Appendix*, Table S6, Fig. S22), which indicated that heterogeneity in *Isodon* diversification rate was triggered by hidden states rather than growth form or habitat. The HiSSE result was corroborated by the FiSSE (Fast, intuitive SSE) model approach that the difference between the speciation rates of herbs (*λ*_0_ = 0.498) and shrubs (*λ*_1_ = 0.524), wet lineages (*λ*_0_ = 0.518) and dry lineages (*λ*_1_ = 0.507), was insignificant (*p* = 0.578 and 0.475, respectively).

For the interactive models, the character-independent MuHiSSE (multistate HiSSE) model with four hidden states was selected as the best-fit model (*SI Appendix*, Table S7), indicating that the interaction between growth form and habitat had no direct effect on *Isodon* speciation rates.

### Niche Preference and Differentiation of *Isodon* Species

Ancestral reconstructions of all 39 environmental variables for herbs and shrubs revealed that factors associated with aridity and precipitation strongly influenced the distinction between the two growth forms (Fig. 3, *SI Appendix*, Fig. S23). Specifically, shrubs were more closely associated with dry and open habitats with less precipitation, higher elevation, lower aridity index, higher soil pH, and herbaceous vegetation, whereas herbs were more frequently found in wetter and more shady habitats.

**Fig. 3.**
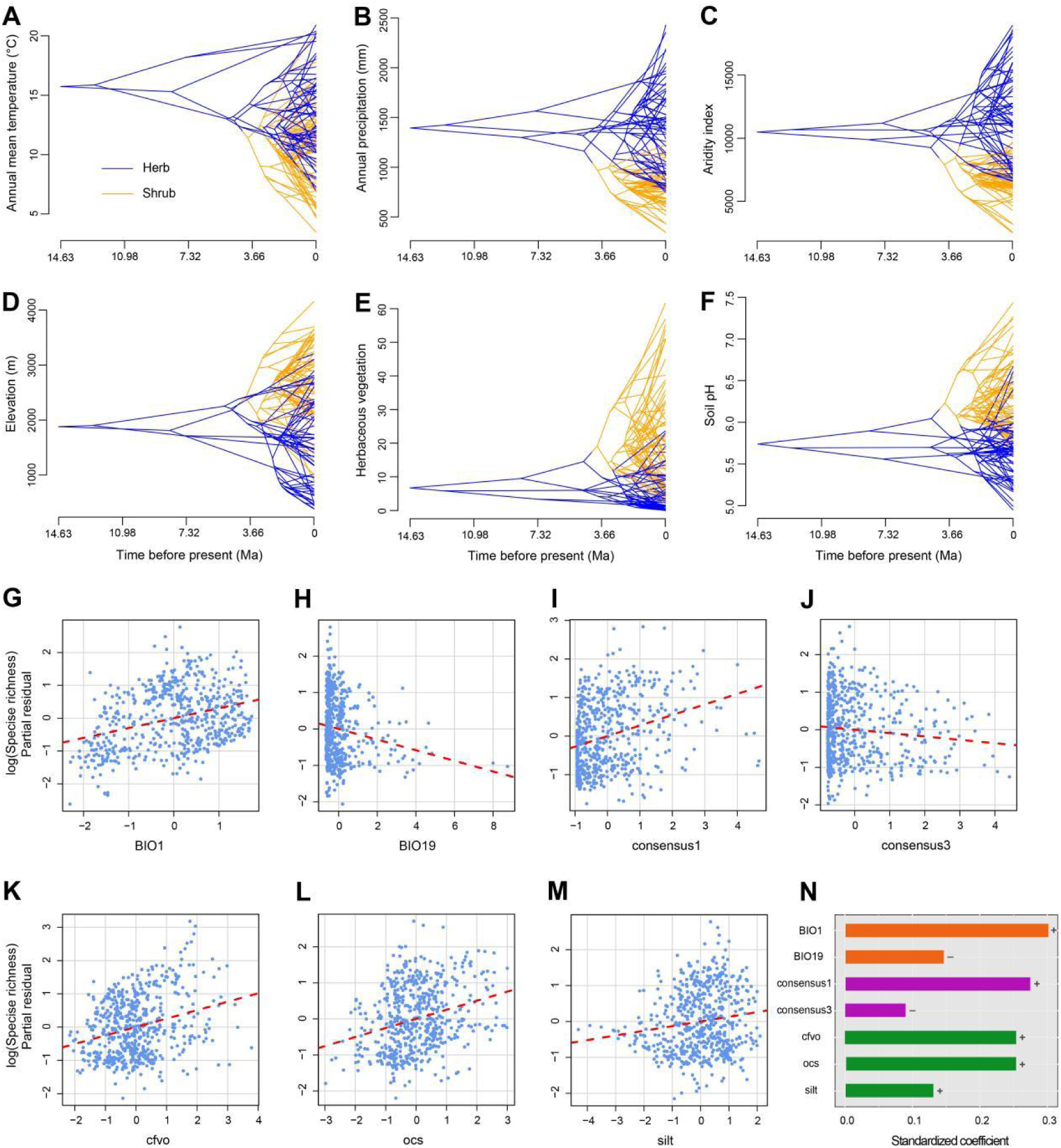
Reconstruction of ancestral niches for the herbs and shrubs and determinants of species diversity of *Isodon*. (A–F) Traitgrams showing the inferred evolution of environmental variables based on the dated phylogeny of *Isodon* in a space defined by the phenotype (y axis). All six variables show distinct differences between the two growth forms according to the t-test (*p* < 0.001). (G–M) Relationships between predictor variables and species richness based on the global multipredictor model across grid cells of 100 km × 100 km. (N) Relative importance of three types of variables (climate, landcover, and soil) in explaining species richness. The positive and negative correlations are indicated as + and −, respectively. Abbreviations and explanations of predictor variables refer to *SI Appendix*, Table S8.

The distribution of grid cell mean species diversity of *Isodon* showed strong geographical variation with the highest diversity in the Hengduan Mountains (*SI Appendix*, Fig. S24). According to the multiple regression analysis, a total of seven environmental variables were significantly correlated with the species richness of *Isodon* (*R^2^*= 0.24) (*SI Appendix*, Tables S8 and S9). Five variables, including annual mean temperature (BIO1), volumetric fraction of coarse fragments (cfvo), organic carbon stocks (ocs), proportion of silt particles (silt), and evergreen/deciduous needleleaf trees (consensus1), were positively correlated with species richness, while the other two variables, including precipitation of coldest quarter (BIO19) and deciduous broadleaf trees (consensus3), were negatively correlated with species richness. Among these variables, soil (especially volumetric fraction of coarse fragments) played the most important role, followed by landcover variables, temperature, and precipitation (Fig. 3, *SI Appendix*, Table S9).

## Discussion

### Rapid Evolutionary Radiation of *Isodon* in the Dry Valleys

RNA sequencing has been shown to be highly efficient for phylogenetic reconstruction at the species level, even for taxa that have undergone a rapid radiation (36, 45–49). This is certainly the case with *Isodon*, for which we have reconstructed a well-resolved phylogeny which includes nearly 90% of extant species and is based on transcriptome sequences with complementary data from genome resequencing (Fig. 1, *SI Appendix*, Figs. S3 and S4). Consistent with previous molecular phylogenetic studies (42, 44), our results suggest that all the sections and series proposed by ref. 39 within *Isodon* are non-monophyletic and that instead, the genus can be divided into four clades (Clade I−Clade IV). However, in previous studies, resolving the phylogenetic relationships within the largest group, Clade IV, has been difficult (42, 44). By utilizing 7,204 orthologs, we are able to resolve four strongly supported clades (Clade IVa−Clade IVd) within Clade IV (Fig. 1, *SI Appendix*, Figs. S3 and S4). Detailed discussion of relationships within each clade is available in *SI Appendix*, *Additional Discussion*.

The crown age of *Isodon* was estimated to be 19.61 Ma (95% HPD: 14.66−26.44 Ma) by ref. 42, i.e., in the early Miocene, somewhat older than the age 14.63 Ma (95% HPD: 10.44−19.35 Ma) estimated here (*SI Appendix*, Fig. S10). This disparity might be caused by the use of the early–middle Oligocene fruit fossil of *Melissa* L. by ref. 42, whose identity was questioned by ref. 50. Though the ancestral range of *Isodon* is unclear, dry valley species first emerged in the Himalaya or the Hengduan Mountains, and a major vicariance event between these two regions occurred later at the crown of clade IVb-IVd (3.91 Ma, 95% HPD: 2.86−5.1 Ma) during the Pliocene (Fig. 2). The acceleration in net diversification rate revealed by our results from BAMM, CoMET, and RPANDA (Fig. 2, *SI Appendix*, Figs. S14 and S15), the shift from herbaceous to shrubby growth form (*SI Appendix*, Fig. S18), and the shift from wet to dry habitat (*SI Appendix*, Figs. S18−S20) are all hypothesized to have occurred on the branch subtending clade IVb-IVd. This striking congruence suggests a rapid evolutionary radiation of shrubby *Isodon* species concentrated in the dry valleys in southwest China during the Pliocene (H1).

Beyond the recent rapid radiation of *Isodon*, our results also suggest a correlated evolution between shrubby growth form and dry valley habitat (H1; *SI Appendix*, Figs. S18, S21). But unlike the parallel evolution of insular woodiness and montane perenniality in triggering high rates of diversification in oceanic island and tropical alpine island-like systems (1, 33), our trait-dependent analyses reveal that a shrubby growth form or dry valley habitat did not promote the diversification rates of *Isodon* (H2; *SI Appendix*, Fig. S22).

Despite a lack of differential diversification rates between different growth forms or habitat types, our findings suggest that aridity and precipitation are major factors differentiating the ecological niche preferences of herbs and shrubs and influencing the distribution patterns of *Isodon* (H3). In particular, shrubs are more accustomed to dry and open habitats, whereas herbs are more frequently found in wet and shady places (Fig. 3, *SI Appendix*, Fig. S23). Moreover, reversals from a shrubby to herbaceous habit in *Isodon* are usually associated with transitions to more closed and humid environments in the Sino-Japanese Floristic Region (Fig. 2, *SI Appendix*, Fig. S18). The shrubby growth form is a trait related to resilience to drier conditions and likely acted as a preadaptation by which *Isodon* species could overcome their humid environmental ancestral state dependence and successfully invaded the seasonally dry river valleys. In addition to the lignified wood cell walls of stems that can avoid or reduce drought-induced air blockage of water transport from roots to leaves (xylem embolism) (51, 52), shrubby species of *Isodon* are diminutive and possess small, coriaceous, and/or densely tomentose leaves to reduce water evaporation (Fig. 1). These drought-adapted characters are shared by many other lineages from dry valleys (5, 7). Even though some herbaceous species also occur in these habitats, they generally grow in more specialized shady places such as in pine forests.

Our multiple regression analyses of the 39 environmental factors suggest that beyond a dry habitat preference, *Isodon* species may also prefer warm needleleaf forests with gravelly soil with large organic carbon stocks. All these related factors may well explain why the highest *Isodon* species richness is sustained in the warm needleleaf forests in southwest China, especially the forests dominated by *Pinus yunnanensis* Frach. from central and northwest Yunnan Province to southwest Sichuan Province (*SI Appendix*, Fig. S24) (53, 54). These pine forests are often distributed in the middle to low elevations of a mountain that except for shrubs, many herbaceous species of *Isodon* from the Hengduan Mountains can also be found here.

Aridity and precipitation as major factors limiting the distribution of *Isodon* species in different growth forms is further supported by our finding that the aridity index is decisive for the rate of niche evolution of *Isodon* (*SI Appendix*, Fig. S17). However, the rate shift position of niche evolution, which is located at the crown of Clade IVd with the rates decelerated sharply (Fig. 2, *SI Appendix*, Fig. S16), was decoupled from that of the diversification rate (Fig. 2), indicating that niche divergence may not be a primary contributor to the rapid radiation of *Isodon*. Nevertheless, the decline of diversification rate from the late Pliocene to present seems to be closely related to the change of rate of adaptation during this time.

Although both growth form and habitat transformation did not trigger the accelerated diversification of *Isodon*, the shrubby growth form may have provided a one-dimensional advantage in the dry valley landscape that allows for the invasion of *Isodon* species in a new adaptive zone. The combination of low dispersal ability, short distances of overall gene flow, and the strong environmental heterogeneity in southwest China could be what drove the rapid diversification of *Isodon* in dry valleys subsequently, i.e., a signature of geographic speciation (55). The diaspores of *Isodon* are nutlets, which are mostly of a small size and a smooth and glabrous surface (44). For species inhabiting seasonally dry habitats, dispersal appears to occur without specialized adaptations. Dry nutlets are shed from the calyx during the dry season and are subject to the uncertainties of such habitats, where rain wash, air currents, or sudden inundations can facilitate the transport of seeds, accompanied by other debris, short distances away from the parent plant (56). The limited seed dispersal of *Isodon* species may restrict gene flow and facilitate rapid speciation at small spatial scales in extensive montane regions with intricate topographic relief and natural barriers, such as tall mountains or areas with suboptimal moisture levels. The high mountains and deep valleys in southwest China provides numerous microhabitats with altitudinal and aspect heterogeneity afforded by a spatially extensive, complex, and highly dissected landscape (57). The long and deep valleys may be conducive to within-valley differentiation of species where the narrowness of the valley constricts gene flow, as in many situations, *Isodon* species are narrowly endemic to small areas with very few co-occurring in any one area, and endemic species restricted to the same valley are usually recovered in the same clade. This is also supported by our BSM analyses that most of the biogeographic events happened in *Isodon* correspond to narrow sympatric speciation, i.e., *in situ* diversification (*SI Appendix*, Table S2).

High-elevation areas in continental mountain ranges are often referred to as “sky islands”, as they are geographically isolated, e.g., by the deep river valleys of southwest China, with limited dispersal but explosive radiation and high endemism, just like oceanic islands isolated by the sea (26, 58–61). From these points of view and exemplified by the rapid radiation of *Isodon*, the deep river valleys of southwest China may show a similar island-like distribution with strong geographical isolation of different valley systems that they could be regarded as “islands of dry valleys” (different from but analogous to ref. 58’s “islands of the upper air”) separated by high mountains. These valleys are also very similar to the Andean valley systems, e.g., the Marañón valley in the Andes, in that both are highly geographically isolated and rich with endemics (62–65).

### East Asian Monsoon and the Assembly of Dry Valley Biomes

As discussed above, the evolution of a shrubby growth form may have permitted *Isodon* species to inhabit seasonally dry habitats, thus facilitating the rapid radiation in the dry valleys in southwest China. The question then arises: was the emergence of dry valleys the catalyst for ecological opportunity with *Isodon*?

Previous work attributed the rapid radiation of *Isodon* to one of the major uplifts of the Tibetan Plateau and subsequent aridification events (42). However, it is becoming clear that such a recent and uniform rise of the Tibetan Plateau never occurred. A complex mountainous landscape with uplands over 4 km in elevation has existed in the Tibetan area throughout the Cenozoic (57). Until the Oligocene, what is now the Tibetan Plateau hosted a deep E-W oriented Central Valley bounded to the north by the Qiangtang uplands and to the south by the Gangdese arc mountains (14). Beginning in the early Eocene (ca. 50 Ma) eastern Tibet underwent rapid uplift, achieving its near present elevation (ca. 3.8 km) by the late Eocene (ca. 35 Ma). Initially this young highland was relatively flat, but the uplift also generated a regional increase in rainfall and a bimodal seasonally wet/dry climate (66, 67). This led to the initiation of drainage systems (68) that today are represented by the major rivers incising the Hengduan Mountains and Yunnan (14). The floor of the Central Tibetan Valley rose through the Eocene and Oligocene and by the early Miocene a near-modern plateau had formed (69, 70).

The Himalaya is the most recent part of the THH region to be uplifted, with the high Himalaya exceeding the height of the Gangdese (and the newly-formed plateau) only becoming established after the middle Miocene (71, 72). Subsequent Himalayan uplift imposed a rain shadow effect exacerbating the drying process of the modern plateau already induced by global cooling (71). The dry valleys in the YZR might have been formed during the middle Miocene and Pliocene as the early Miocene temperate woodlands of the central plateau were replaced by grasslands and shrubs (73). Our biogeographic results also suggested that the shrubby ancestors of *Isodon* had emerged in the dry valleys in the YZR during the Pliocene (Fig. 2).

As the diversity center of *Isodon*, the dry valleys in the Hengduan Mountains accommodate most of the woody species. However, the timing of the formation of these dry valleys is complicated and controversial, since only limited studies are available and there is no agreement on when the deeply incised river valleys formed (74). The Hengduan mountains are today the most species-rich area of the THH region (75), and several quantitative palaeoaltimetry studies have demonstrated that the Hengduan Mountains (including central Yunnan, see ref. 48) achieved near-present elevations by the end of Paleogene (66, 67, 76, 77). Though some isostatic rebound (uplift) continued in places into the late Miocene (78), the pronounced topological relief of the Hengduan Mountains was undoubtedly shaped by the large-scale river incision (79–81) and increased erosion during a period of intensified monsoon precipitation in the middle Miocene (74). The scale of the erosion is evidenced by enhanced rock cooling rates during this period (ca. 13−9 Ma) (79, 82).

Recent paleoclimate modelling studies have revealed that the growth of the Tibetan topography, especially the orographic change to the present-day northern Tibetan Plateau, significantly altered the climate of East Asia as the East Asian monsoon developed and increased overall precipitation (83–85). Regional precipitation increased steadily throughout the Cenozoic and started a sharp intensification since the middle Miocene, reaching peak intensity during ca. 12−4 Ma in the late Miocene and Pliocene (“super monsoon”) (83). Notably, the shift of the net diversification rate of *Isodon* is coincidental with the end of the “super monsoon” (Fig. 2). Using a paleoenvironment-dependent diversification model, we also find that the diversification rate of *Isodon* is highly negatively correlated with the strength of the East Asian monsoon (*SI Appendix*, Fig. S15). Taken together, we may infer that increased precipitation enhanced river downcutting, particularly during the Miocene, shaping the deep river valleys. Following erosional removal of load, local relief was further enhanced into the Pliocene by isostatic rebound, further downcutting and thus deeper valleys. Moderate clockwise rotation of eastern Tibet may have begun in the Eocene (86) but at ca. 10 Ma further clockwise rotation about the eastern Himalayan syntaxis (86, 87) began and subsequently aligned the ridges between the drainage systems in the southern Hengduan Mountains into the N-S direction they are today. These ridges would have interacted with the predominantly eastward flowing summer monsoon winds and generated strong local rain shadow effects within the valleys during the Pliocene. Decreased precipitation due to the weakening of monsoon and the rain shadow effect cast by the highly elevated mountains during the Pliocene probably contributed to the aridification and formation of the dry valleys in the Hengduan Mountains.

The evolution of the monsoon system has significantly affected the vegetation in East Asia, resulting in a transition from arid/semi-arid to humid/semi-humid dominated vegetation (84). The intensification of the East Asian monsoon has also been suggested to have promoted species diversification of many plant lineages within East Asia (3, 36, 84, 88, 89). Our study further demonstrates that the interplay between topography and changes in the dynamics of the East Asian monsoon since the middle Miocene might have also shaped the formation of the dry valleys in southwest China and promoted the differentiation of microhabitats along elevational gradients and slope aspects, and thus triggered the ecological opportunity for *Isodon* species and their adaptive radiation during the Pliocene.

In contrast to the recent configuration of dry valley biome in southwest China and rapidly diversified and species-rich clades in this biome as revealed by our study, the seasonally dry tropical forests of the deeply entrenched inter-Andean valleys are occupied by old, slowly diversifying, and species-poor clades (62, 64, 90). This distinction might be partially explained by the geological histories and assembly of flora of the THH region and the Andes. Though the THH region harbors the world’s most species-rich temperate alpine flora, extant alpine lineages within this area can be traced back to the early Oligocene in the Hengduan Mountains, with an increased diversification promoted by mountain building and intensification of the Asian monsoon during the early to middle Miocene (3). The Páramo, the high elevation Andean grassland biome and the world’s fastest evolving biodiversity hotspot, on the contrary, is of recent origin and characterized by clades that have diversified very rapidly in the Pliocene and Pleistocene, favored by the fast uplift rates between 8 and 5 Ma in the Northern Andes (1, 65). Seasonally dry Andean forests at lower elevations show the opposite pattern in that they have been present for nearly 60 million years and assembled gradually over the past 20 Ma (65).

Our study contributes to a deeper understanding of the evolutionary dynamics and ecological and biological drivers influencing the flora of the unique and complex landscape in southwest China. Comparative studies of additional lineages exhibiting both dry valley and non-dry valley diversity are needed to extend and generalize the insights of our findings.

## Materials and Methods

### Taxon Sampling and Sequencing

An ongoing comprehensive taxonomic revision indicates that the genus *Isodon* comprises ca. 140 species, of which 23 species remain undescribed or require new name combinations. We therefore treated these species as *Isodon* sp. 1–23 in this study. We assembled a comprehensive sampling of *Isodon* by including 171 accessions of 126 taxa (including 123 species and three varieties) from all major geographic distribution regions of the genus, representing all recognized sections and series of ref. 39 and all four clades recovered in ref. 42 and ref. 44. We selected nine species from six genera of five subtribes of tribe Ocimeae as outgroups.

Except for the whole genome sequence of one accession of *I. rubescens* that was retrieved from the China National GeneBank DataBase (CNGBdb) (accession no. CNP0002852; ref. 91), we generated all sequences for this study, including transcriptomes for 140 ingroup accessions and nine outgroup accessions, as well as genome resequencing data for 30 ingroup accessions (*SI Appendix*, *Additional Methods*). Voucher information for all samples included in present study is available in Data S1.

### Data Processing and Orthology Inference

Most species of *Isodon* were shown to be diploids except for the two African endemics which were inferred to be of allopolyploid origin (42, 92). Therefore, we followed the “phylogenomic dataset construction” pipeline in https://bitbucket.org/yanglab/phylogenomic_dataset_construction/ (93) and the method used by ref. 94 for transcriptome read processing, assembly, translation, and orthology inference. These methods were designed for complex scenarios of polyploidy and reticulate evolution from various datasets. We briefly describe major procedures below, with more details in the *SI Appendix*, *Additional Methods*. After correcting sequencing errors and removing adapters and low-quality bases for raw reads, we *de novo* assembled processed reads with Trinity v.2.8.5 (95) and removed low-quality and chimeric transcripts following ref. 96. We clustered filtered transcripts into putative genes with Corset v.1.07 (97) and only retained the longest transcript of each putative gene (98). Finally, we translated the filtered transcripts with TransDecoder v.5.0.2 (99) and removed identical CDS.

For the genome resequencing data, we first processed raw reads using fastp v.0.12.4 (100). We then carried out assembly of nuclear loci in HybPiper v.2.1.3 (101) using the CDS of the genome of *I. rubescens* as references after running the “check_targetfile” and “fix_targetfile” commands in Hybpiper. We flagged loci with potential paralogs when multiple contigs cover at least 75% of the reference sequence length and then we extracted each locus together with flagged paralogs using the “paralog_retriever” command in HybPiper.

We combined all CDS assembled from the 149 transcriptomes and 30 resequenced individuals, as well as the genome of *I. rubescens* for downstream orthology inference. We carried out homology inference on CDS using reciprocal BLASTN, followed by orthology inference using the “monophyletic outgroup” approach from ref. 93, keeping only ortholog groups with at least 90 tips. We then re-aligned the new fasta files written from ortholog trees using PRANK v.170427 (102), removed aligned columns with more than 70% missing data using Phyx (103), and retained alignments with at least 300 characters and 90 accessions. Using this method, in total we identified 7,204 orthologous genes.

### Phylogenetic Analyses

We performed phylogenomic analyses using maximum likelihood (ML) with a supermatrix implemented in IQ-TREE v.2.0.3 (104), and a multispecies coalescent model with ASTRAL-III v.5.7.8 (105). For the ML analysis, we identified the best-fit substitution model for the concatenated 7,204 orthologous dataset via ModelFinder (106) implemented in IQ-TREE, and determined node support with 1,000 UFBS replicates (107). To estimate a multispecies coalescent tree, we first inferred 7,204 individual gene trees using RAXML v.8.2.12 (108) with a GTRGAMMA model, and 100 BS replicates to assess clade support. After collapsing the nodes with less than 10% BS support value using the “nw_ed” function of Newick utilities v.1.6 (109), we used individual gene trees to estimate a species tree in ASTRAL with LPP (110) to assess clade support.

We visualized discordance among the ML tree and the coalescent tree using an R script retrieved from ref. 36. To distinguish strong conflict from weakly supported branches, we evaluated tree conflict and branch support with Quartet Sampling (QS) which subsamples quartets from an input tree and concatenated alignment to assess the confidence, consistency, and informativeness of each internal branch using the relative frequency of the three possible quartet topologies at each node (111). We used the concatenated dataset and the coalescent tree as input, with the number of replicate quartet searches per branch set to 500 and the log-likelihood cut-off to 2. We then visualized the QS results using an R script (https://github.com/ShuiyinLIU/QS_visualization). We also calculated the quartet support for the coalescent tree and two alternative topologies and carried out a polytomy test for all branches of the coalescent tree using the parameters “-t 8” and “-t 10” in ASTRAL, respectively (112). A *p*-value of <0.05 is considered to reject the null hypothesis of a polytomy.

As the two African species were hypothesized to be of allopolyploid origin based on a previous study, and their phylogenetic placements showed strong topological conflict between the nuclear and plastid trees (42, 44), we further tested the hybrid origin of the two species and other potential reticulation events among major clades of *Isodon* using species network searches with the command “InferNetwork_MPL” in PhyloNet v.3.8.2 (113), with the individual gene trees as input. Due to computational considerations, we reduced our taxon sampling to one outgroup and 13 ingroup taxa, including 1–3 representative species from each of the major clades in *Isodon*. We pruned the individual gene trees to include one outgroup (*Siphocranion macranthum*) and at least six of the 13 ingroups. We carried out three network searches by allowing one to three reticulation events and five independent runs for each search. To estimate the optimum number of reticulations, we optimized the branch lengths and inheritance probabilities, and computed the likelihood of the best scored network from each of the three maximum reticulation events searches. We used the command “CalGTProb” in PhyloNet (114) to estimate the network likelihoods given the individual gene trees. We also inferred potential reticulation events within Clade I by selecting 11 ingroup taxa from this clade, and one outgroup from Clade III (*I. ternifolius*), with the remaining settings the same as above.

### Divergence Time Estimation

We employed two methods, a Bayesian analysis using BEAST v.1.10.4 (115) and the penalized likelihood method implemented in treePL (116), to estimate the divergence time of *Isodon*. Given the computational resources needed for Bayesian methods of large datasets, we used the SortaDate package (117) to filter the top 30 “best” loci (i.e., using a gene shopping approach). This package determines which gene trees are most clock-like, have the least topological conflict with the coalescent tree, and have informative branch lengths. We used the GTR+G model for site substitutions, the birth–death model for the tree prior, and the uncorrelated relaxed lognormal model for the clock prior. We fixed the tree topology using the coalescent tree inferred using ASTRAL. Within Ocimeae, only the putative early Eocene pollen fossils of *Ocimum* L. identified by ref. 118 are available, but previous phylogenetic studies of Lamiaceae used these fossils as a constraint for the crown node of Nepetoideae. Therefore, we used the 95% HPD from the most recent Lamiaceae-wide dating analysis (119) as a constraint for the crown node of the Ocimeae clade using a normal distribution, with a mean of 41.89 and a standard deviation of 5.56 (95% HPD: 31−52.77 Ma). We also constrained the crown age of the clade excluding *Siphocranion* Kudô and *Lavandula* L. using a normal distribution prior, with a mean of 23.01 and a standard deviation of 4.66 (95% HPD: 13.88−32.13 Ma; ref. 119). We ran the BEAST analyses for 200 million generations, logging parameters every 1,000 generations.We checked the stationary state and convergence in Tracer v.1.7 (120) to ensure that all parameters had effective sample sizes (ESS) above 200. Then we generated the maximum clade credibility (MCC) tree with median age and 95% HPD for each node using TreeAnnotator 1.10.4 (of the BEAST package) with the first 10% of trees discarded as burn-in. For the treePL analysis, because the coalescent species tree does not have terminal branch lengths, we initially optimized the branch lengths in RAxML using the “-f e” option to yield branch lengths in expected per-site substitutions. We employed the same calibrations for treePL as those used in the BEAST analyses but with hard minimum and maximum age constraints. Following determination of best optimization parameters in the prime step, we conducted a cross-validation analysis to choose the best smoothing value, i.e., the one with the lowest chi-square value in the resulting cvoutfile. Subsequently, we dated the tree using the best optimization parameters and the best smoothing value selected above.

### Occurrence Data and Environmental Variables

To collect the distribution information of *Isodon* for further biogeographic and ecological niche analyses, we downloaded occurrence data of all *Isodon* species from GBIF (the global biodiversity information facility, https://www.gbif.org/) and added specimen records from 10 herbaria (BM, E, IBSC, K, KUN, KYO, L, P, PE, TI; abbreviations follow ref. 121) and our own field expeditions. We carefully assessed these datasets and removed erroneous records (i.e., occurrences in the ocean, ice sheets, deserts, and urban areas), duplicates, and cultivated records. Finally, we assembled a total of 6,708 unique distribution records for 140 species, including the 126 species sampled in the phylogenomic analysis (Data S2; *SI Appendix*, Fig. S1).

We collected 39 environmental variables for these records, including 19 bioclimatic variables and one topographical layer (elevation) (ref. 122; https://worldclim.org/data/worldclim21.html), 11 soil variables within 30 cm of soil horizon (ref. 123; https://files.isric.org/soilgrids/latest/data_aggregated/1000m/), six landcover classes (ref. 124; https://www.earthenv.org/landcover) as well as aridity index and annual potential evapotranspiration (ref. 125; https://figshare.com/articles/dataset/Global_Aridity_Index_and_Potential_Evapotranspiration_ET0_Climate_Database_v2/7504448/6) (*SI Appendix*, Table S8; Data S2). All variables were at a resolution of 30 arc seconds. We used the mean values of the variables for each species in the subsequent analyses of niche rate estimation and ancestral niche reconstruction.

### Ancestral Area Estimation

We carried out ancestral area reconstructions using the R package BioGeoBEARS v.1.1.1 (126) as implemented in RASP 4.2 (127). Based on the extant distribution patterns of *Isodon* species, we delimited six geographical areas: (A) the Himalaya, together with part of the Hajar Mountains in the Arabian Peninsula; (B) the Hengduan Mountains and Yunnan Plateau; (C) subtropical China (central, southern, and eastern China, but excluding the tropical parts of southern China), part of northern China and the Russian Far East, and the Korean Peninsula; (D) South and Southeast Asia; (E) the Japanese Archipelago; and (F) Africa. We ran the analyses on the MCC chronogram from the BEAST analysis but with outgroups and multiple accessions of species pruned. We set the maximum number of ancestral areas at each node to four as *Isodon* species are never found in more than four areas. As the founder-event speciation (j) parameter remains controversial (128, 129), we only tested three non-nested models, including DEC (dispersal–extinction–cladogenesis; ref. 130), DIVA-like (a likelihood implementation of dispersal–vicariance analysis; ref. 131), and BayArea-like (a likelihood implementation of Bayesian inference of historical biogeography for discrete areas). We treated the model with the highest weighted AIC corrected for sample size (AICc_wt) as the best model. To estimate the number and type of biogeographical events, we also conducted BSM implemented in BioGeoBEARS (132). We estimated event frequencies by taking the mean and standard deviation of event counts from 50 BSMs.

### Macroevolutionary Rate Estimation

We used three different approaches to investigate the temporal dynamics of diversification rates of *Isodon* (H1). First, we employed BAMM v.2.5.0 (133) to estimate diversification rates through time and detect potential rate shift across the *Isodon* phylogeny. We specified the globalSamplingFraction as 0.90 and estimated prior values for BAMM using the setBAMMpriors function in the R package BAMMtools v.2.1.10 (134). We carried out the BAMM analysis using the MCC tree from the BEAST analysis but with outgroups and multiple accessions of species pruned. We ran the Markov chain Monte Carlo (MCMC) for 100 million generations and sampled every 100,000 generations. After assessing convergence using the R package CODA v.0.19.4 (135) to ensure ESS above 200, we discarded the initial 20% samples of the MCMC run as burn-in and used BAMMtools to plot the best shift configuration with the maximum a posteriori probability and diversification rates through time. Second, we applied the CoMET model (136) as a complement to BAMM to jointly estimate time-continuous rates and detect abrupt changes in speciation and extinction rates. We implemented CoMET in the R package TESS v.2.12 (137) and ran MCMC for 10 million generations with default priors and not allowing for mass extinctions. Third, to explore the effects of global paleotemperature and East Asian monsoons on the diversification of *Isodon*, we also used the R package RPANDA v.2.2 (138) to fit a series of time-, temperature-, and monsoon-dependent likelihood diversification models. We obtained the temperature data from the Cenozoic temperature curves documented and revised by ref. 139 and ref. 140 and acquired the East Asian monsoon data using the hematite/goethite proxy, assessed through measurements at 565 and 435 nm wavelengths within the color spectra of a freshly cut core retrieved from Ocean Drilling Program Site 1148 in the South China Sea (141). We modelled speciation and extinction dependencies as all possible combinations of constant and exponential relationships as well as pure-birth models. We used the equations *λ*(E) = *λ*_0_ × e*^α^*^E^ and *μ*(E) = *μ*_0_ × e*^β^*^E^ for modelling exponential dependence, where *λ*_0_ and *μ*_0_ represent the speciation and extinction rates for a specified environmental variable, respectively. The parameters *α* and *β* denote the rates of change in speciation and extinction relative to the environment, with positive values indicating a positive effect of the environment on speciation or extinction, and vice versa (142). We compared the likelihood support values and chose the model with the lowest AICc value as the best diversification model. We randomly sampled 200 trees from the BEAST posterior distribution with outgroups and multiple accessions of species removed to accommodate dating uncertainties. We used the R package PSPLINE v.1.0 (143) to visualize variation in speciation rates with paleoenvironmental variables.

To estimate rates of niche evolution in *Isodon*, we first ordinated all environmental variables using phylogenetic principal component analysis (PCA) implemented in the R package phytools v2.1-1 with the “phyl.pca” function (144). Then, we conducted complementary runs using the BAMM trait model on the first axis of the phylogenetic PCA of niche data. We ran the MCMC for 100 million generations and sampled every 100,000 generations. We discarded the initial 20% samples of the MCMC run as burn-in and used BAMMtools to plot the rates through time.

### Ancestral State Reconstruction and Correlated Evolution Test

To test if rate shifts of *Isodon* are associated with shifts to a shrubby growth form and evolutionary transitions to dry valleys (H1), we reconstructed the evolutionary history of both characters (growth form and habitat type). We compiled information on growth forms from literature (39, 40, 145) and our own field observation. Species of *Isodon* are either perennial herbs or shrubs, but continuous variation in wood development in the aboveground stem among species often makes the boundary between herbaceousness and woodiness fuzzy (146, 147). For example, the transitional growth form “undershrubs” or “subshrubs” are used in the literature for some *Isodon* species (e.g., *I. ternifolius*). These species are mainly herbaceous but woody at the base. As detailed wood anatomical data for the genus are not available, and given that we were only interested in understanding the shift toward increased wood formation, we divided growth form of *Isodon* into herb and shrub, and categorized species that develop a wood cylinder extending toward the upper parts of the stems which are usually persistent in winter as shrubs, and those species whose stems are not woody or only woody at the base, often withering in winter, as herbs. We classified species into two habitat types based on rainfall and aridity regimes: dry (valley) and wet. We defined dry valley species as those with a mean annual precipitation < 1,200 mm (148) and an average aridity index < 10,000. We then treated the remaining species as wet lineages. We reconstructed the ancestral state of growth form and habitat of *Isodon* using ML and the Markov k-state one-parameter model of evolution for discrete unordered characters in Mesquite v.3.81 (149). We also reconstructed the ancestral niche of *Isodon* using the first axis of the phylogenetic PCA of niche data and conducted the analysis using the “contMap” function in the phytools package with model choice via AICc; the model set included BM, Ornstein-Uhlenbeck (OU), and early-burst (EB) models.

To assess the correlated evolution between dry valleys and shrubby growth form (H1), we initially examined the level of phylogenetic signal for these two categorical traits following ref. 150. We carried out the phylogenetic signal tests using the pruned MCC tree and the *D* statistic (151), as implemented in the R package caper v1.0.3 (152). The *D* value ranges from 0 to 1, where *D* = 0 indicates that the trait evolves following a random model (lacking phylogenetic signal). Conversely, *D* = 0 suggests that the trait evolution adheres to the Brownian model, while *D* < 0 signifies a strong phylogenetic signal, indicating minimal observed differences between sister clades for the given binary trait (151). We determined significance in our estimate of *D* with 1,000 permutations. We then performed correlated evolution tests using Pagel’s correlation method (153) for binary characters states implemented in phytools package. We tested both independent and dependent models. For a dependent model approach, we tested three models in which the rates of change in habitat type (dependent variable) were dependent on the state of the growth form and/or the rates of change in growth form were dependent on the state of the habitat type.

### Trait-Dependent Analyses

We investigated whether two traits, growth form and habitat, were associated with increased rates of diversification in *Isodon* (H2). We first examined correlation using the HiSSE model in the R package hisse v.2.1.11 (154). The HiSSE model is an extension to optimize the binary state speciation and extinction model (BiSSE), which estimates turnover (*τ* = speciation (*λ*) + extinction (*µ*)) rates, extinction fraction (*ε* = *μ*/*λ*), and transition rates (*q*) associated with a binary character, while accounting for the estimation of unobserved traits that could affect diversification rates. We fitted 24 models to the *Isodon* phylogeny that differed in how their parameters were constrained. Detailed model description are available in *SI Appendix*, *Additional Methods*. We also used the nonparametric FiSSE model (155) as a complementary analysis for measuring the robustness of our HiSSE results by calculating speciation rates for the binary traits and assessing statistical differences between the two states.

To evaluate if speciation rates of dry valley shrubs are highest (H2), we built interactive models encompassing both characters (growth form and habitat type) and three combined states (i.e. herbs from wet habitat, dry valley herbs, dry valley shrubs). We used the multistate speciation and extinction with hidden traits – MuHiSSE (156) approach and fitted eight different models (*SI Appendix*, *Additional Methods*). We ran all models in the hisse package and assessed the best-fit model based on the AICc values.

### Niche Preference and Differentiation of *Isodon* Species

To test if shrubby species exhibit broader drought tolerance compared to herbaceous species (H3), we compared the niche preferences of species with different growth forms by reconstructing the ancestral state for each of the 39 environmental variables using the “fastAnc” function and visualizing the degree of similarity of each variable among species with different growth forms using the “make.simmap” and “phenogram” functions in the phytools package. We also compared the mean within groups to those of the whole sample, using a Welch two sample t-test. To do so, we tested the null hypothesis of no difference between groups with *p* = 0.05, after assuring for variance homogeneity and normal data distribution.

We then tested if drought-related factors are major driving forces of the species richness of *Isodon* (H3). We first calculated the species richness by matching the distribution data of all 140 species of *Isodon* to the 100 km × 100 km grid cells of the entire land of the world projected using Behrmann Equal-Area Cylindrical in ArcGIS v10.8 (Data S3). We then extracted all 39 environmental variables as mean values within each 100 km × 100 km grid cell (except for elevation which was extracted as range within each grid cell) (Data S3). To reduce collinearity among variables, we reduced the initial set of 39 environmental variables to ten variables with weak pairwise correlations (|r| < 0.5). We built a multiple regression model (ordinary least squares, OLS) for all *Isodon* species, with species richness as the response variable and ten environmental variables as predictors. Additional details of the methods are available in *SI Appendix*, *Additional Methods*.

## Acknowledgments

We would like to thank Dr. Jian Liu, Dr. Hang-Hui Kong, Dr. Yong-Sheng Chen, Dr. Lu Sun, and Dr. Diego F. Morales-Briones for their help in data analyses. Thanks are also extended to Prof. Colin E. Hughes and Dr. Richard Ree for their constructive comments on earlier drafts of this manuscript and to the staff of the Germplasm Bank of Wild Species in Southwest China for providing leaf material and collection information. This work was supported by the Ten Thousand Talents Program of Yunnan (Grant No. YNWR-QNBJ-2018-279) and the open research project of the Germplasm Bank of Wild Species, Kunming Institute of Botany, Chinese Academy of Sciences to CLX, the Yunnan Fundamental Research Projects (Grant Nos. 202101AT070159, 202301AT070303, 202101AU070067) and the Yunnan Revitalization Talent Support Program “Young Talent” Project to YPC, the Science and Engineering Research Board of the Government of India (Grant No. CRG/2018/003499) to PS, and the CAS Interdisciplinary Team of “Light of West China” and Yunnan Revitalization Talent Support Program “Innovation Team” Project to CLX.

