## Supplementary Information for "Rapid radiation of a plant lineage sheds light on the assembly of dry valley biomes"

### Additional Methods

#### Taxon Sampling and Sequencing

An ongoing comprehensive taxonomic revision indicates that the genus *Isodon* comprises ca. 140 species, of which 23 species remain undescribed or require new name combinations. We therefore treated these species as *Isodon* sp. 1–23 in this study. We sampled a total of 171 accessions of 126 taxa (including 123 species and three varieties) from all major regions across the geographic distribution of *Isodon* in Asia and Africa, and from major river valleys in southwest China (Figs. S1, S2), representing all recognized sections and series of ref. 1 and all four clades recovered in ref. 2 and ref. 3. We selected nine species from six genera of five subtribes of tribe Ocimeae as outgroups.

Except for the whole genome sequence of one accession of *I. rubescens* (Hemsl.) H. Hara that was retrieved from the China National GeneBank DataBase (CNGBdb) (accession no. CNP0002852; ref. 4), we generated all sequences for this study, including transcriptomes for 140 ingroup accessions and nine outgroup accessions, as well as genome resequencing data for 30 ingroup accessions. Voucher information for all samples included in present study is available in Data S1.

For transcriptome sequencing, we extracted total RNA from young and healthy leaves and shoots, which were collected and frozen in liquid nitrogen before storage at –80 °C using the RNAprep Pure Plant Kit (Tiangen Biotech, Beijing, China). We sent the RNA to the Novogene Corporation (Tianjin, China) for preparation of cDNA libraries and sequencing on an Illumina NovaSeq 6000 instrument, generating 150 bp paired-end reads and ca. 6 Gb raw data for each sample.

For genome resequencing, we extracted total genomic DNA either from silica-gel-dried leaves using a modified CTAB method (5) or from herbarium material using the DNeasy Plant Mini Kit (Tiangen Biotech, Beijing, China). We sent the DNA to BGI Genomics (Shenzhen, China) for library preparation and sequencing on an Illumina HiSeq 2000 instrument, generating 150 bp paired-end reads and ca. 20 Gb raw data for each sample.

#### Data Processing and Orthology Inference

Most species of *Isodon* were shown to be diploids except for the two African endemics which were inferred to be of allopolyploid origin (2, 6). Therefore, we followed the "phylogenomic dataset construction" pipeline in https://bitbucket.org/yanglab/phylogenomic_dataset_construction/ (7) and the method used by ref. 8 for transcriptome read processing, assembly, translation, and orthology inference. These methods were designed for complex scenarios of polyploidy and reticulate evolution from various datasets. We first corrected sequencing errors in raw reads with Rcorrector (9) and removed adapters and low-quality bases using Trimmomatic v.0.36 (10). We then *de novo* assembled processed nuclear reads with Trinity v.2.8.5 (11) using default settings and removed low-quality and chimeric transcripts following the approach of ref. 12, using *I. rubescens* as the reference proteome (4). We clustered filtered transcripts into putative genes with Corset v.1.07 (13) and only retained the longest transcript of each putative gene (14). We translated the filtered transcripts with TransDecoder v.5.0.2 (15) with default settings and used proteomes of *Arabidopsis thaliana* (L.) Heynh. and *I. rubescens* to identify open reading frames. Finally, we reduced the coding sequences (CDS) from translated amino acids with CD-HIT v.4.7 (16) to remove redundancy (-c 0.99).

For the genome resequencing data, we first removed adaptors and low-quality bases (Phred scores < 20) from raw reads using fastp v.0.12.4 (17). We then carried out assembly of nuclear loci in HybPiper v.2.1.3 (18) using the CDS of the genome of *I. rubescens* as references after running the “check_targetfile” and “fix_targetfile” commands in Hybpiper. We flagged loci with potential paralogs when multiple contigs cover at least 75% of the reference sequence length and then we extracted each locus together with flagged paralogs using the “paralog_retriever” command in HybPiper.

We combined all CDS assembled from the 149 transcriptomes and 30 resequenced individuals, as well as the genome of *I. rubescens* for downstream orthology inference. First, we performed an all-by-all BLASTN search on CDS using an E value cutoff of 10 and max_target_seqs set to 1000. We then filtered raw BLAST output with a hit fraction of 0.4 and clustered putative homolog groups using MCL v.14-137 (19) with a minimum minus log-transformed E value cutoff of 5 and an inflation value of 1.4. We discarded clusters with fewer than 90 tips and aligned the remaining clusters with MAFFT v.7.407 (20) using the accuracy-oriented method E-INS-i and a maximum of 1000 iterative refinement cycles. After removing aligned columns with more than 90% missing data using Phyx (21), we built homolog trees using RAXML v.8.2.12 (22) with the GTRGAMMA model of nucleotide substitution. We pruned tips longer than 10 times the average of its sister, or longer than 0.4 expected substitutions per site. Additionally, we removed mono- and paraphyletic tips that belonged to the same individual by keeping only the tip with the highest number of characters in the trimmed alignment and cut internal branches longer than 0.4 expected substitution per site, keeping all resulting subclades with a minimum of 90 taxa. We carried out homolog tree inference, tip and outlier removal, and a second round of deep paralog cutting using the same settings above (except for an absolute_cutoff of 0.2) to obtain final homologs.

We implemented orthology inference following the “monophyletic outgroup” approach from ref. 7, keeping only ortholog groups with at least 90 tips. To maximize the number of orthologs obtained, we only set the three species of *Siphocranion* Kudô as outgroups for this step. We then re-aligned the new fasta files written from ortholog trees using PRANK v.170427 (23), removed aligned columns with more than 70% missing data using Phyx, and retained alignments with at least 300 characters and 90 accessions. Using this method, in total we identified 7,204 orthologous genes.

**Trait-Dependent Analyses**

We investigated whether two traits, growth form and habitat, were associated with increased rates of diversification in *Isodon* (H2). We first examined correlation using the hidden state speciation and extinction model in the R package hisse v.2.1.11 (HiSSE; ref. 24). The HiSSE model is an extension to optimize the binary state speciation and extinction model (BiSSE), which estimates turnover ($\tau$ = speciation ($\lambda$) + extinction ($\mu$)) rates, extinction fraction ($\varepsilon = \mu/\lambda$), and transition rates ($q$) associated with a binary character, while accounting for the estimation of unobserved traits that could affect diversification rates. We fitted 24 models to the *Isodon* phylogeny that differed in how their parameters were constrained. Two models were BiSSE-like models (i.e., without hidden states), where turnover parameters were free to vary, and extinction fraction or transitions rates were constrained between growth forms states, depending on the model. Four models corresponded to different character-independent diversification (CID) models, which assumed that rate differences are associated with two (CID-2) or four (CID-4) hidden states and extinctions or transitions were constrained between states. The remaining 18 HiSSE models were trait-dependent diversification models with hidden states, different combinations of parameter constraints, and two different types of transition matrices, one restricted (default) and the other allowing all transitions between character states. We assessed the best-fit model based on the AICc values. We also used the nonparametric FiSSE model (Fast, intuitive SSE model) (25) as a complementary analysis for measuring the robustness of our HiSSE results by calculating speciation rates for the binary traits and assessing statistical differences between the two states.

To evaluate if speciation rates of dry valley shrubs are highest (H2), we built interactive models encompassing both characters (growth form and habitat type) and three combined states (i.e. herbs from wet habitat, dry valley herbs, dry valley shrubs). We used the multistate speciation and extinction with hidden traits – MuHiSSE (26) approach, which is a HiSSE extension that handles multiple states. Then, we fitted eight different models. One of these was a null model where diversification rate variation is not caused by character states. Two models were character-dependent diversification without a hidden state (i.e. MuSSE-like model). Another two models were character-dependent diversification with a hidden state and the last three models were character-independent diversification with two to four hidden states. We ran all MuHiSSE models in the hisse package and assessed the best-fit model based on the AICc values.

**Niche Preference and Differentiation of *Isodon* Species**

To test if shrubby species exhibit broader drought tolerance (H3), we compared the niche preferences of species with different growth forms by reconstructing the ancestral state for each of the 39 environmental variables using the "fastAnc" function and visualizing the degree of similarity of each variable among species with different growth forms using the "make.simmap" and "phenogram" functions in the R package phytools v2.1-1 (27). We also compared the mean within groups to those of the whole sample, using a Welch two sample t-test. To do so, we tested the null hypothesis of no difference between groups with $p = 0.05$, after assuring for variance homogeneity and normal data distribution.

We then tested if drought-related factors are major driving forces of the species richness of *Isodon* (H3). We first calculated the species richness by matching the distribution data of all 140 species of *Isodon* to the 100 km × 100 km grid cells of the entire land of the world projected using Behrmann Equal-Area Cylindrical in ArcGIS v10.8 (Data S3). We then extracted all 39 environmental variables as mean values within each 100 km × 100 km grid cell (except for elevation which was extracted as range within each grid cell) (Data S3). To reduce collinearity among variables, we reduced the initial set of 39 environmental variables to ten variables with weak pairwise correlations ($|r| < 0.5$), including three bioclimatic variables (BIO1, annual mean temperature; BIO15, precipitation seasonality; and BIO19, precipitation of coldest quarter), aridity index (AI), three soil variables (cfvo, volumetric fraction of coarse fragments; silt, proportion of silt particles; and ocs, organic carbon stocks), and three landcover variables (consensus1, evergreen/deciduous needleleaf trees; consensus3, deciduous broadleaf trees; and consensus5, shrubs). We built a multiple regression model (ordinary least squares, OLS) for all *Isodon* species, with species richness as the response variable and ten environmental variables as predictors. We log-transformed the response variable and further evaluated the multi-predictor model for multi-collinearity using Variance Inflation Factors (VIF) in the R package car v3.1-2 (28) and removed the predictors with VIFs greater than 5 before model selection. We scaled all predictors to a mean of zero and variance of one before the analysis to make the direct comparison of regression coefficients and the OLS model residuals approximated a normal distribution. We used a stepwise regression based on the AIC to derive a minimum adequate model which has the smallest number of predictors and retains the highest explanatory power. For the minimum adequate model, we obtained the standardized coefficient of each predictor to compare the relative importance of predictors in explaining *Isodon* species richness.

**Additional Discussion**

Consistent with previous molecular phylogenetic studies (2, 3), our results suggest that all the sections and series proposed by ref. 1 within *Isodon* are non-monophyletic and that instead, the genus can be divided into four clades (Clade I−Clade IV) (Fig. 1). All species recovered in Clade I are herbs with reddish-brown glands on all plant parts. The latter character was considered a useful diagnostic synapomorphy of this clade (compared to the colorless glands in the remaining clades) (2, 3, 29). In the study by ref. 3, the placement of *I. scrophularioides* exhibited a large discrepancy being placed in different subclades of Clade I in the plastid and nuclear gene trees. This is consistent with our PhyloNet analyses which suggest that *I. scrophularioides* may be a hybrid between *I. phulchokiensis* and the common ancestor of the *I. oreophilus* ~ *I. flavidus* clade, with a major genomic contribution from *I. phulchokiensis* (*SI Appendix*, Fig. S9). This also explains the overall morphological similarity between *I. phulchokiensis* and *I. scrophularioides*, as well as the short branch length (*SI Appendix*, Figs. S3 and S4) and low quartet support of node A (*SI Appendix*, Fig. S8). The African Clade II is recovered as sister to Clade I in the present study, but was shown to be a sister of clade III-IV in the plastid tree (3). Clade II has been suggested to be of allopolyploid origin by ref. 2, which is further confirmed by our PhyloNet analyses (*SI Appendix*, Fig. S9). With a major genomic contribution from the common ancestor of Clade I, species of Clade II show overall morphological similarity with species of Clade I, but have lost the reddish-brown glands.

We further recognize four strongly supported subclades within Clade IV. All species of Clade IVa were previously treated as a single species, *I. coetsa* (Buch.-Ham. ex D. Don) Kudô, by ref. 1, whereas both molecular and morphological evidence suggest that each of these represent a distinct lineage. Clade IVb comprises all shrubs distributed in the Himalaya (with *I. rugosus* extending to the Hajar Mountains in the Arabian Peninsula), especially the dry valleys of the YZR. Though this clade receives maximum support in both the coalescent tree and the ML tree, its monophyly is challenged by short branch length (*SI Appendix*, Figs. S3 and S4) and QC < -0.05 (*SI Appendix*, Fig. S6). Additionally, the quartet scores for the main topology and two alternative topologies of node B are 0.37, 0.26, and 0.36, respectively (*SI Appendix*, Fig. S8). This conflict may be explained by the possible hybrid origin of *I. pharicus*, which is placed in a subclade within Clade IVb, between *I. rugosus* from another subclade of Clade IVb and the common ancestor of clade IVc-IVd (*SI Appendix*, Fig. S9). Apart from the basal subclade that comprises shrubs, all remaining species of Clade IVc are herbs distributed in East Asia (from the Hengduan Mountains to the Japanese Archipelago). A significant majority of species of *Isodon*, and species with a shrubby habit from the dry valleys in the Hengduan Mountains are placed in Clade IVd, but relationships within this clade exhibit considerable conflict between the coalescent tree and the ML tree (*SI Appendix*, Fig. S5), and many nodes receive low support values or quartets scores (*SI Appendix*, Figs. S3, S4, and S8). These conflicts could be caused by hard polytomies, incomplete lineage sorting, and/or gene flow. More in-depth analyses of the gene tree conflicts are needed to further clarify the complex evolutionary history of Clade IVd.

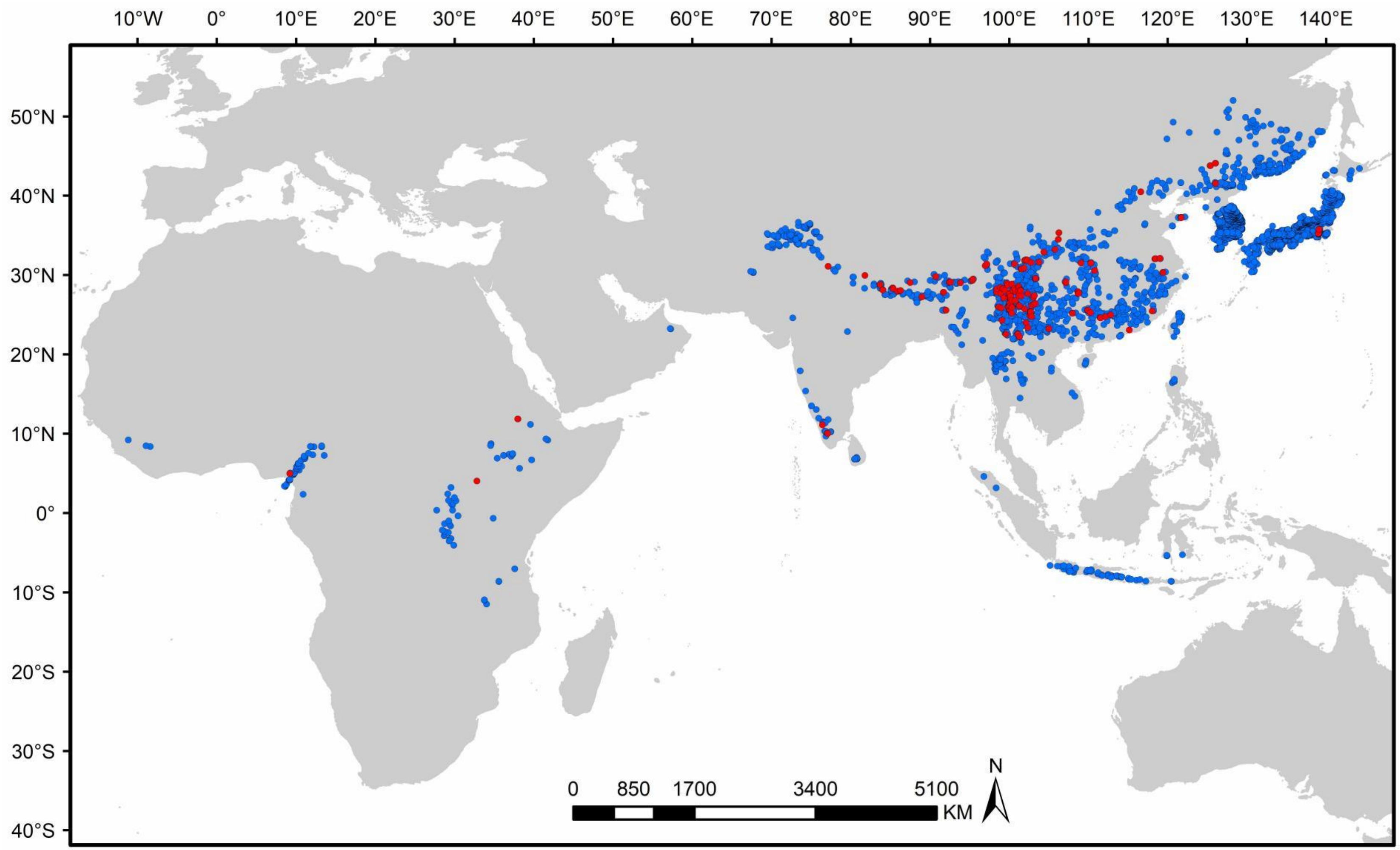


**Fig. S1.** Distribution and sampling map of *Isodon*. Accessions sampled in present study are marked in red (with cultivated vouchers excluded).

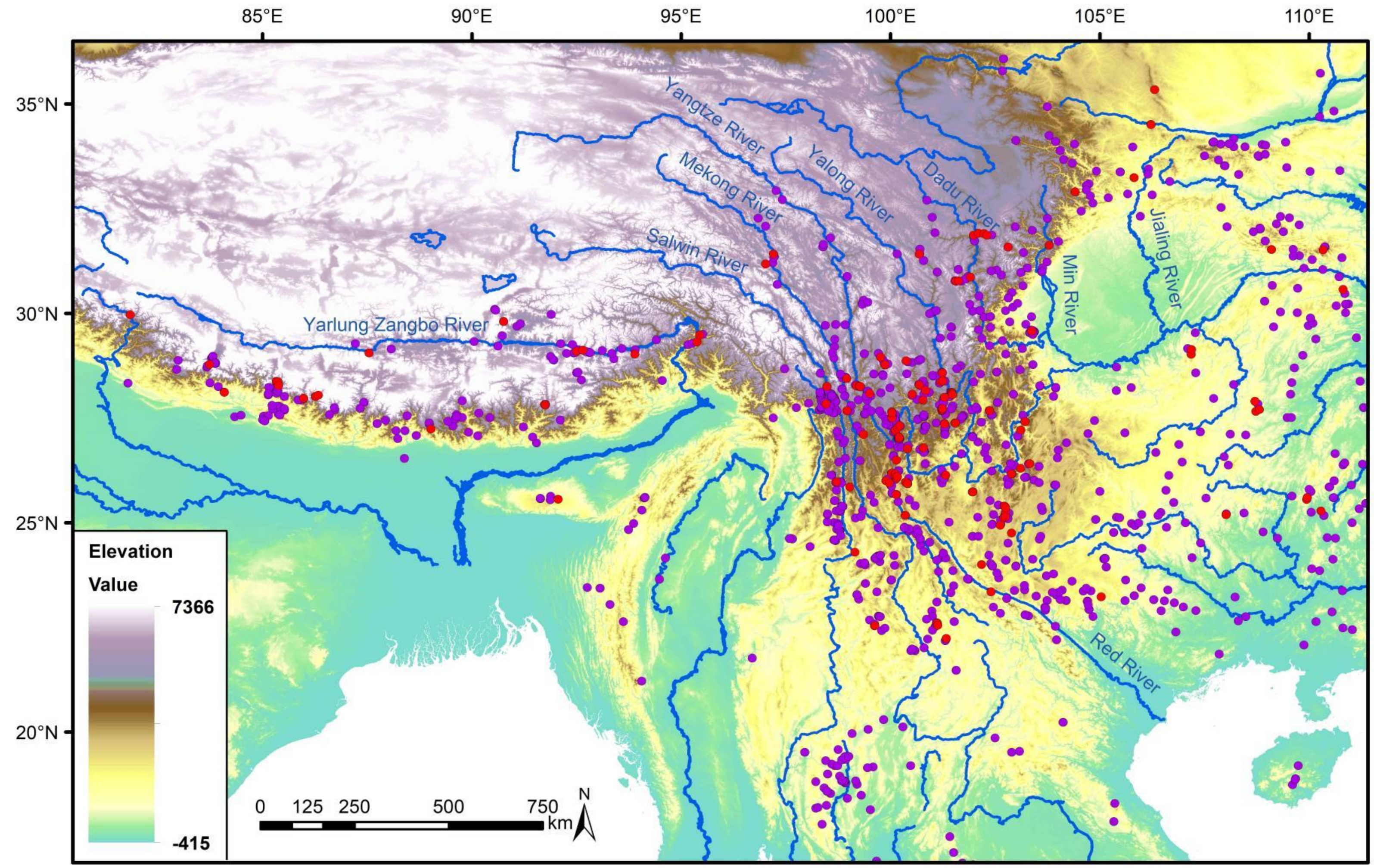


**Fig. S2.** Distribution and sampling of *Isodon* in southwest China. Accessions sampled in present study are marked in red.

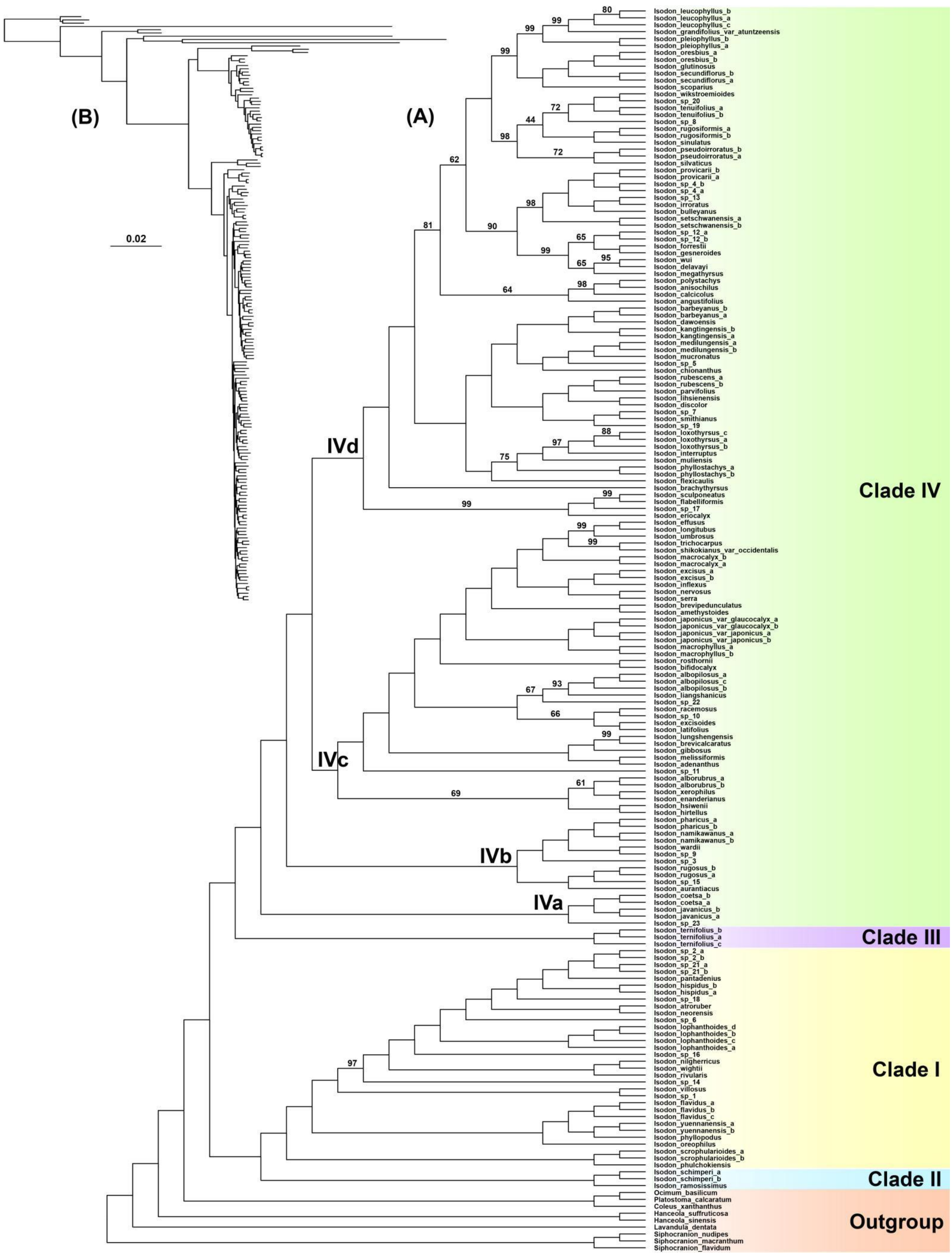


**Fig. S3.** Cladogram (A) and phylogram (B) of the maximum-likelihood tree of *Isodon* estimated using the concatenation method of IQ-TREE based on the dataset of 7,204 low-copy nuclear genes. Ultrafast bootstrap (UFBS) values = 100% are not shown, and numbers on nodes indicate UFBS values < 100%.

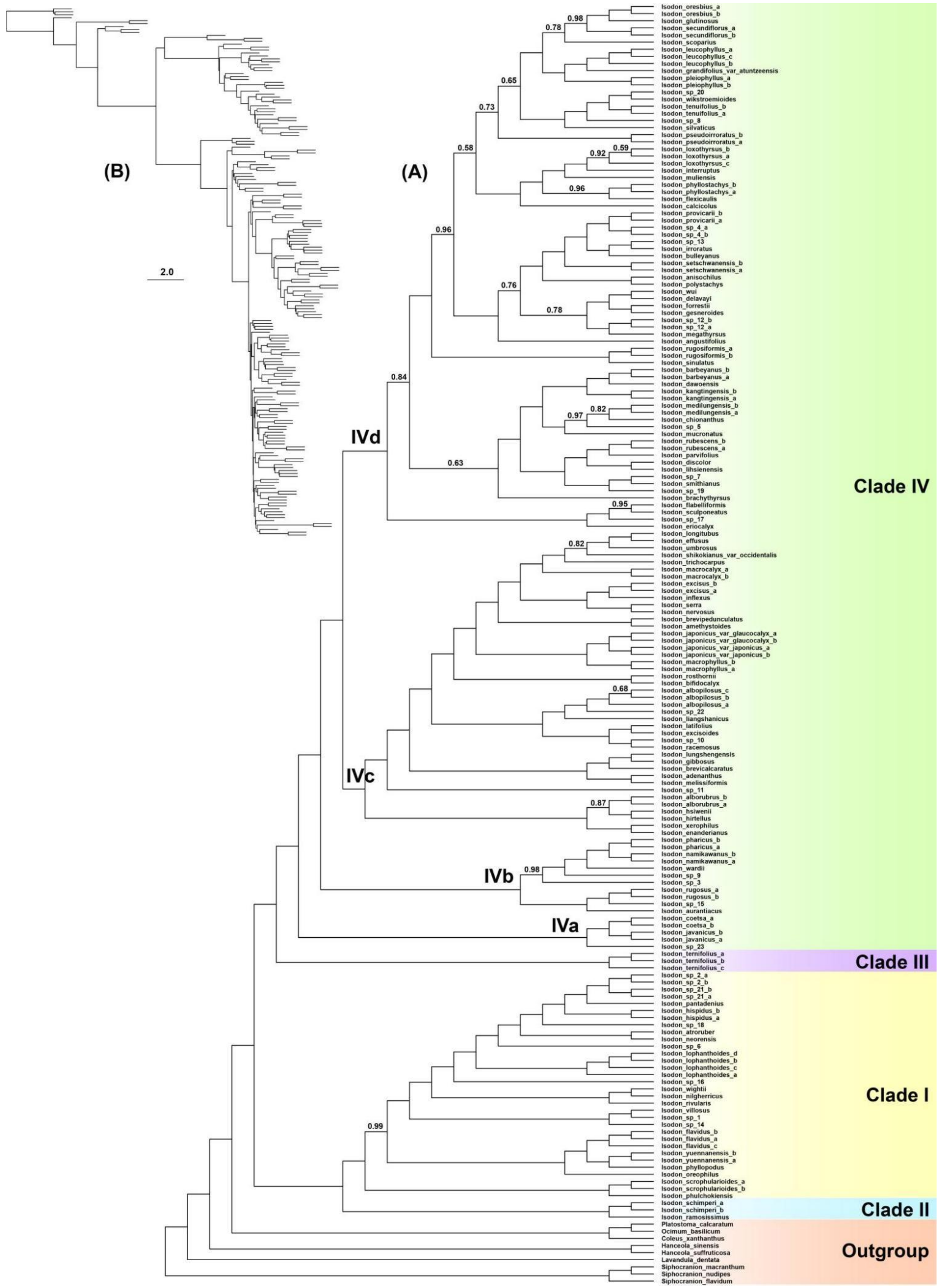


**Fig. S4.** Cladogram (A) and phylogram (B) of the species tree of *Isodon* estimated using the coalescent method of ASTRAL based on the single gene trees inferred from 7,204 low-copy nuclear genes. Local posterior probabilities (LPP) = 1.00 are not shown, and numbers on nodes indicate LPP < 1.00.

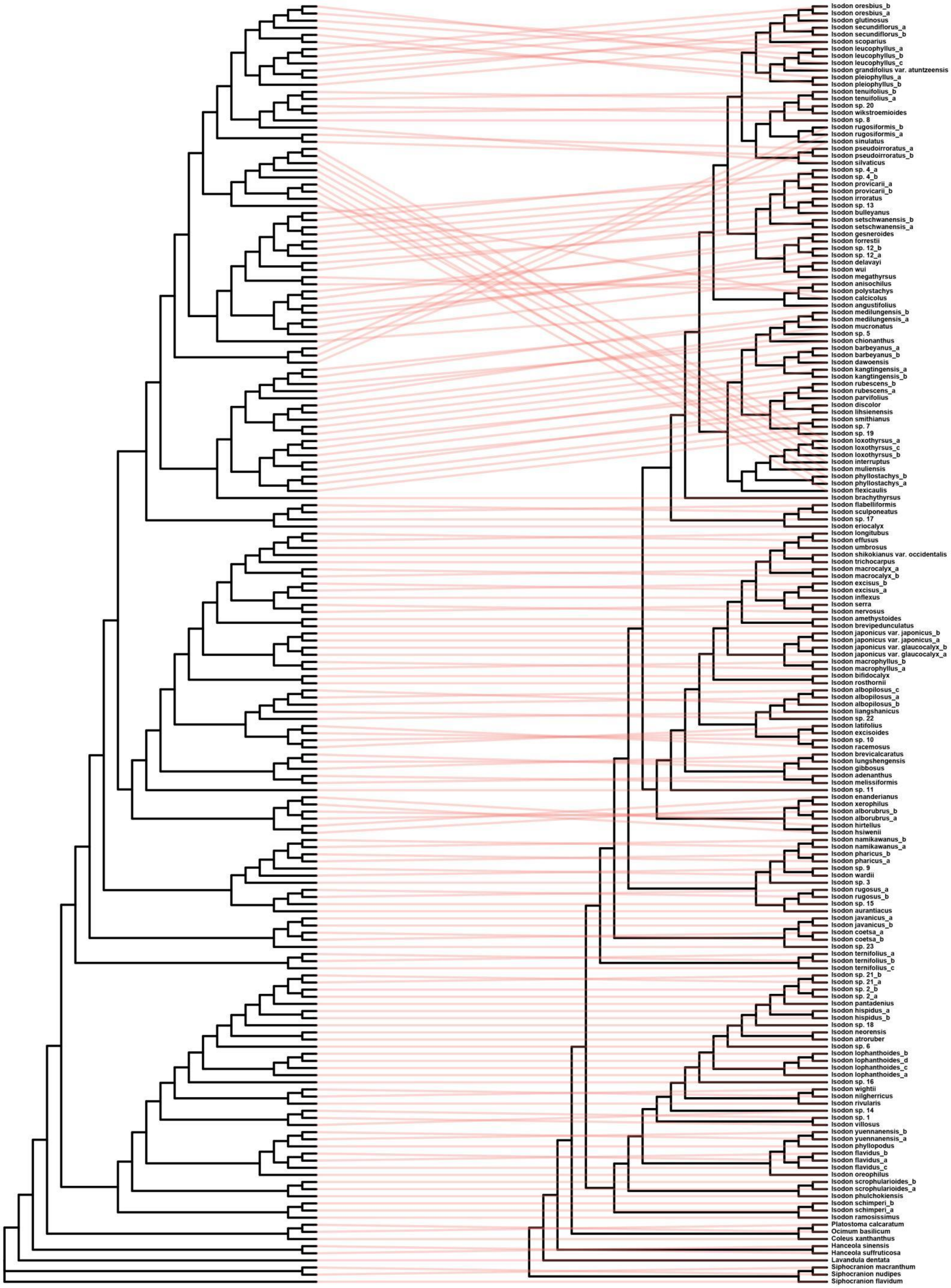


**Fig. S5.** Visualization of discordance between the topologies of the coalescent tree (left) and the maximum-likelihood tree (right) of *Isodon*.

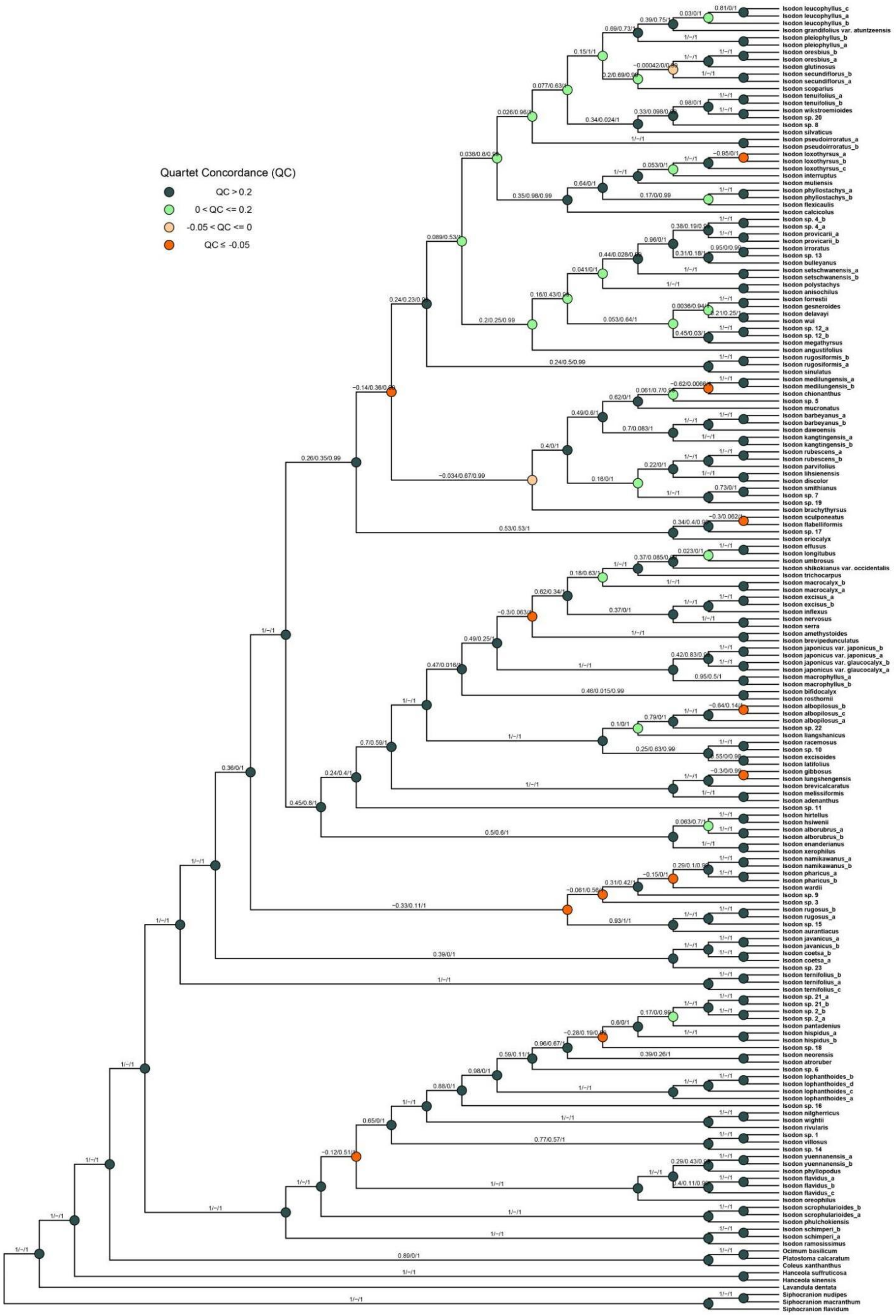


**Fig. S6.** Quartet sampling results of the species tree of *Isodon* estimated using the coalescent method of ASTRAL based on the single gene trees inferred from 7,204 low-copy nuclear genes. Quartet concordance (left), quartet differential (middle), and quartet informativeness scores (right) are given above the branches.

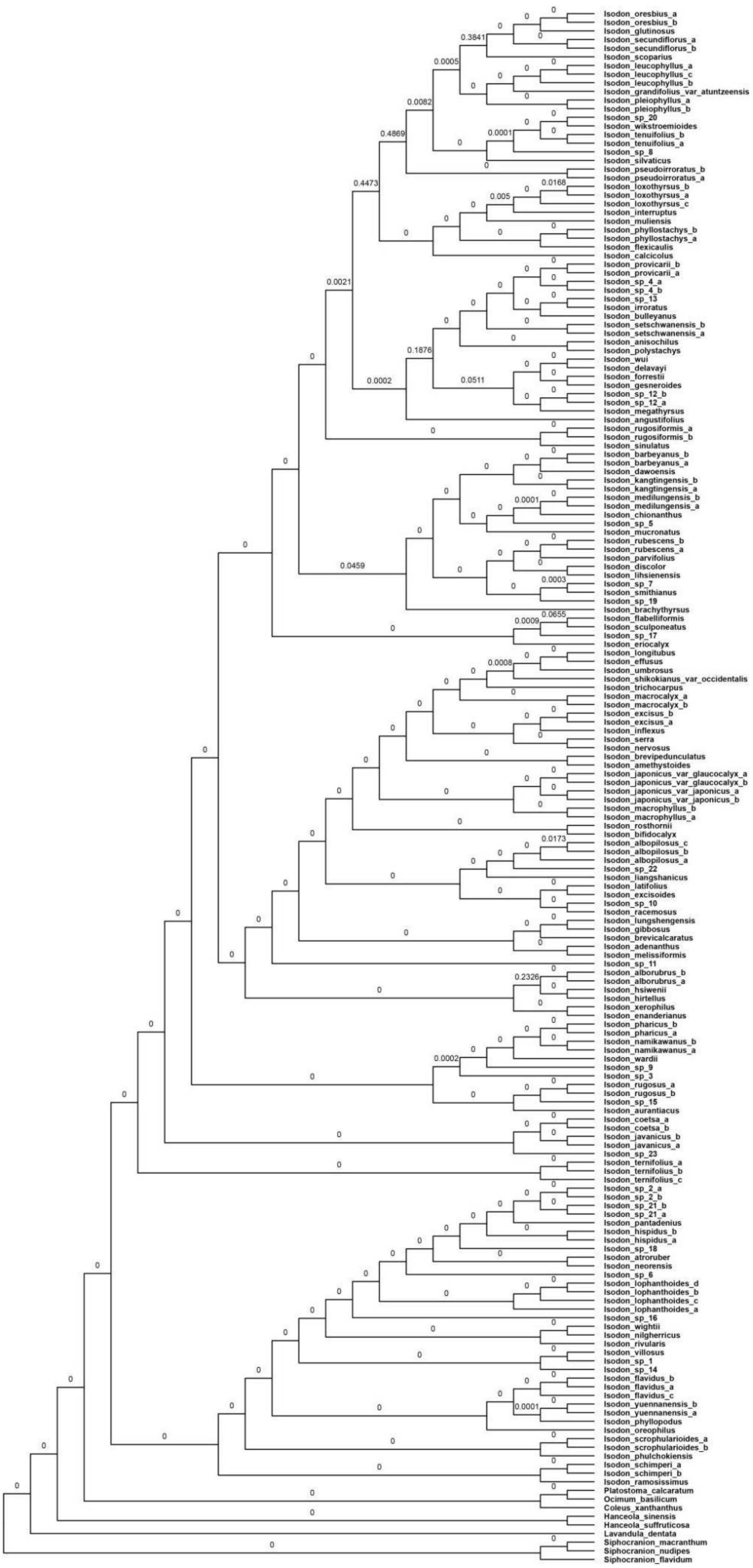

**Fig. S7.** Polytomy test of the species tree of *Isodon* estimated using the coalescent method of ASTRAL based on the single gene trees inferred from 7,204 low-copy nuclear genes, with *p*-value from the polytomy test shown above branches.

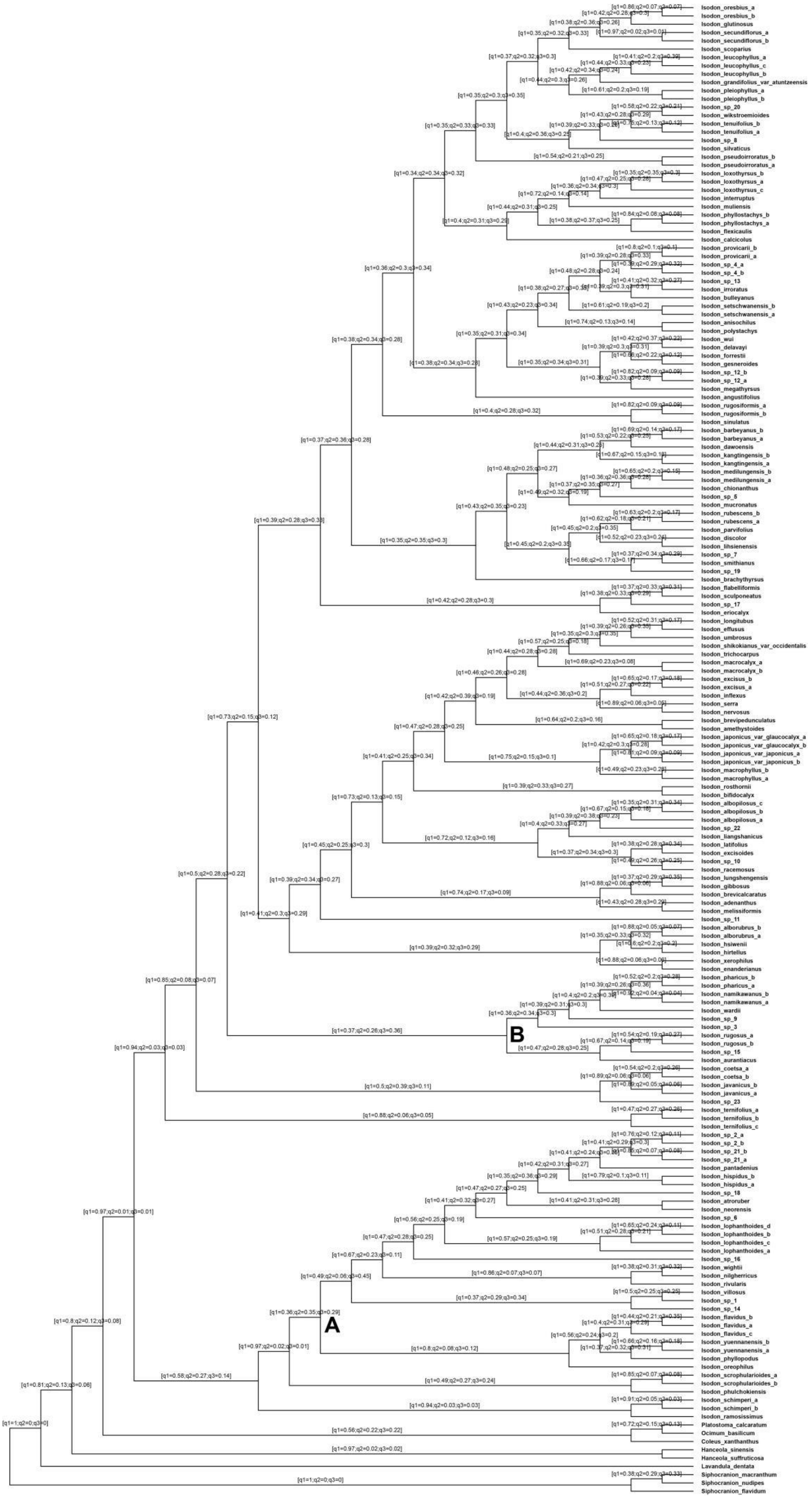


**Fig. S8.** Quartet supports for the species tree of *Isodon* estimated using the coalescent method of ASTRAL based on the single gene trees inferred from 7,204 low-copy nuclear genes and two alternative topologies.

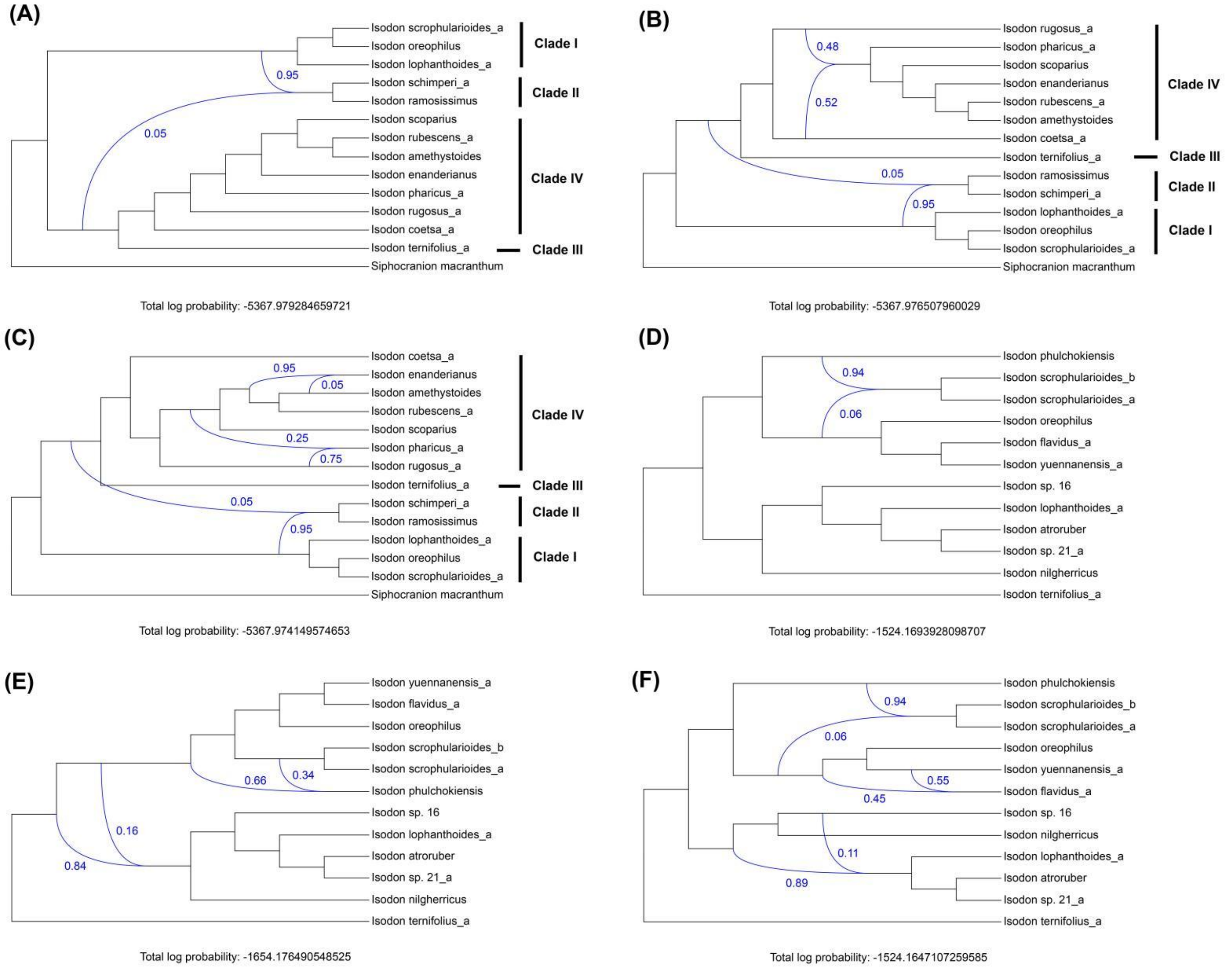


**Fig. S9.** The optimal phylogenetic network of *Isodon* (A–C) and Clade I (D–F) inferred using PhyloNet, with the number of reticulations as 1–3.

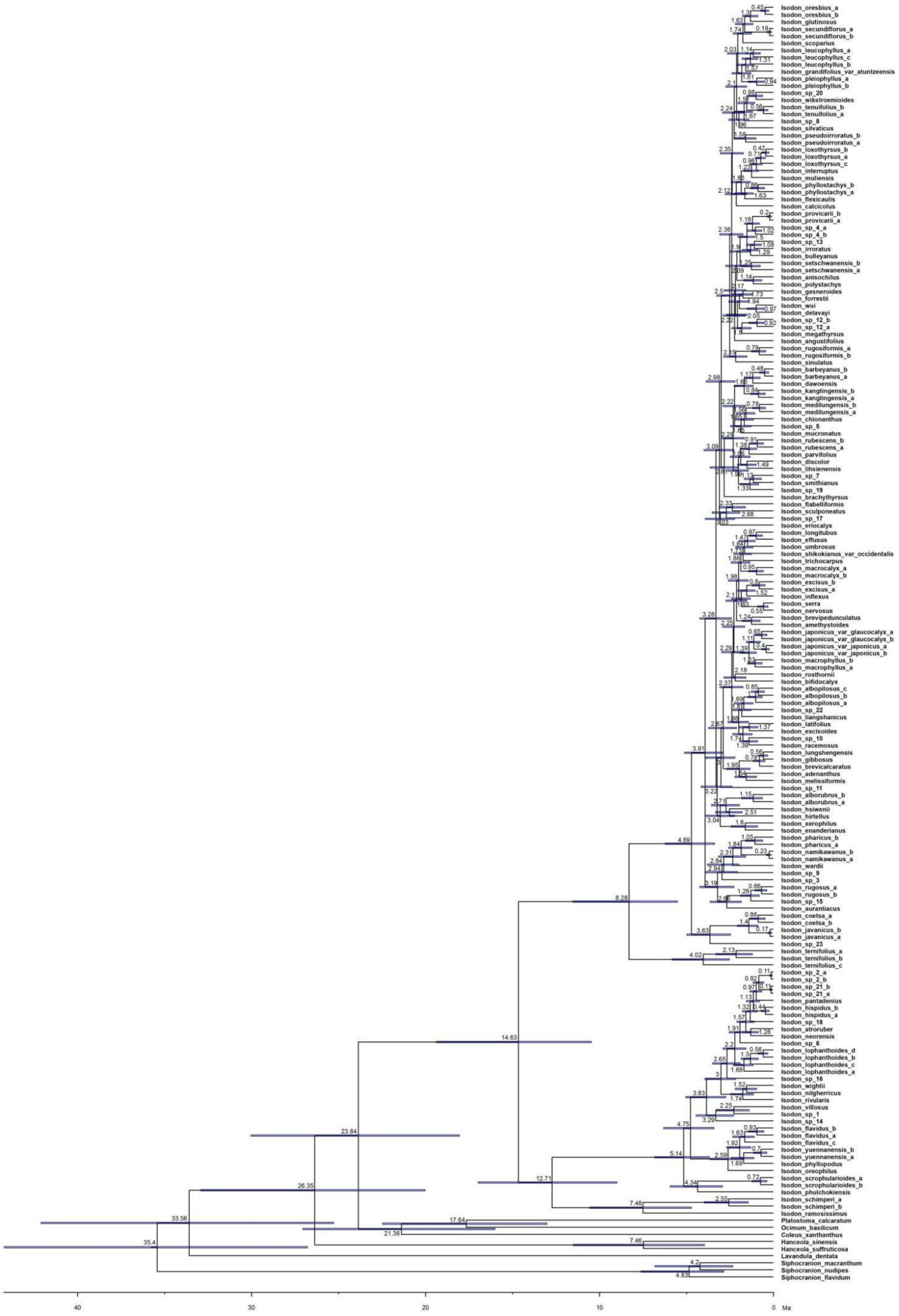


**Fig. S10.** Dated phylogeny of *Isodon* using BEAST. The numbers at nodes indicate median divergence times (Ma). Blue bars represent 95% HPD of node age.

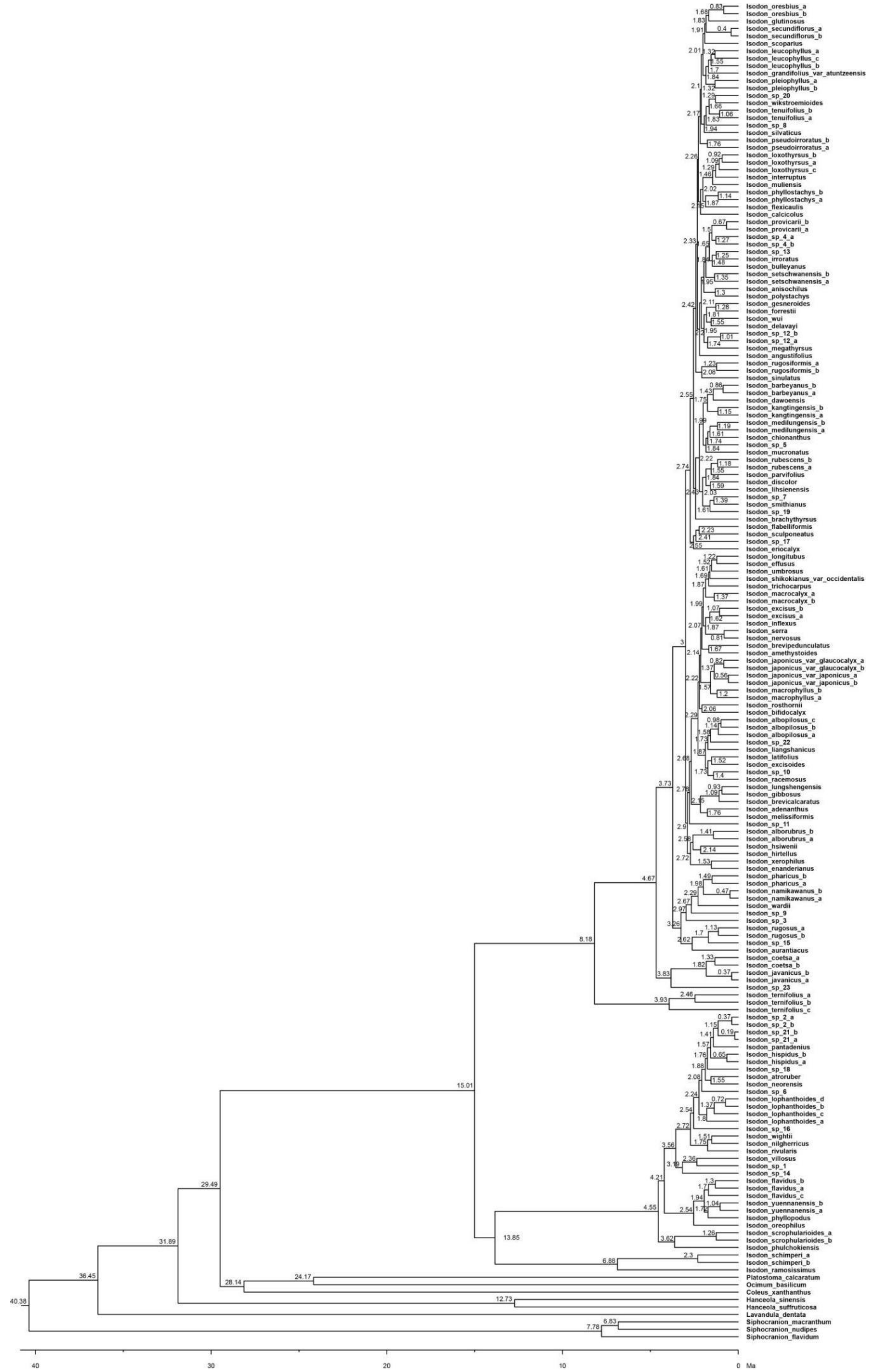


**Fig. S11.** Dated phylogeny of *Isodon* using treePL. The numbers at nodes indicate divergence times (Ma).

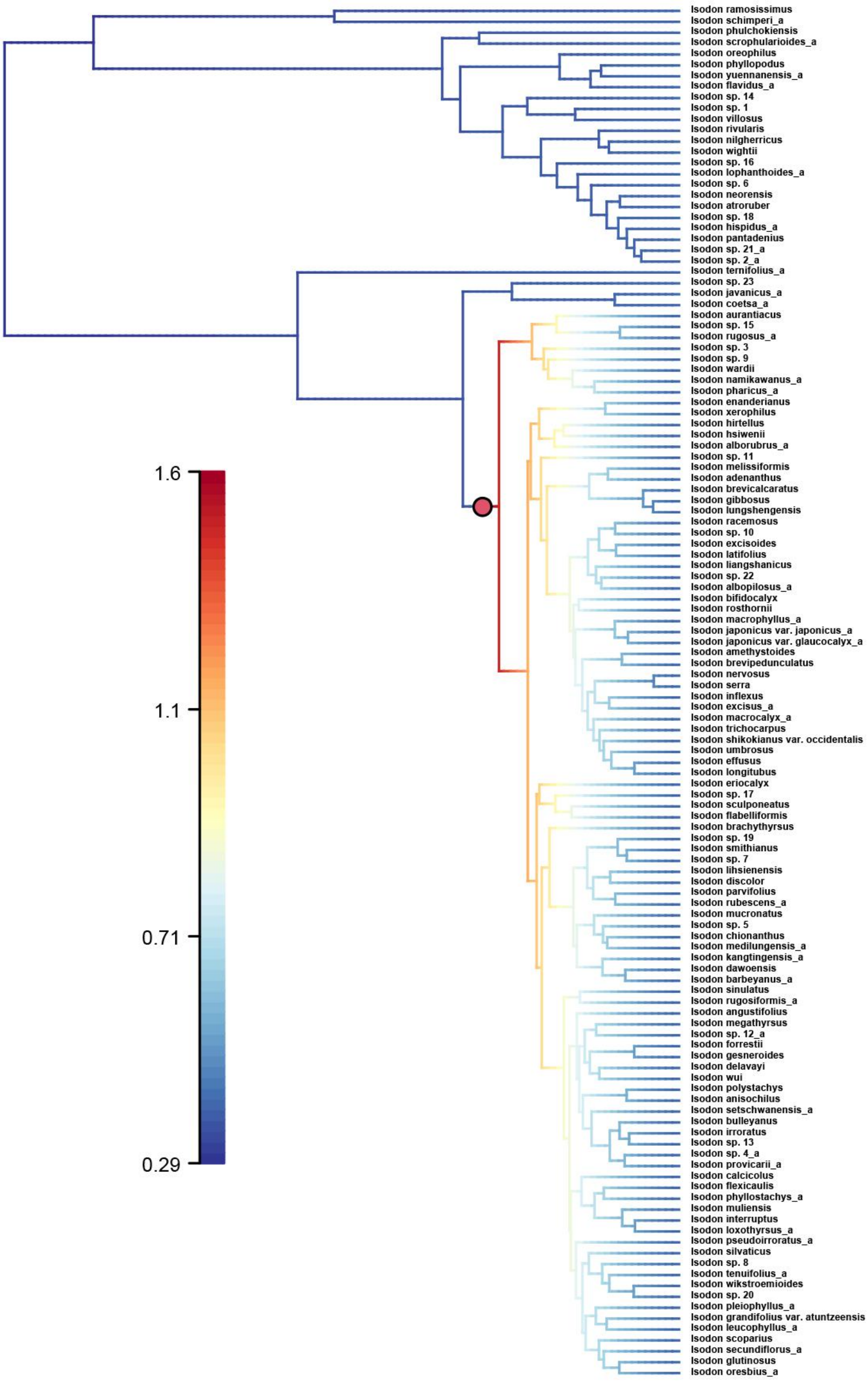


**Fig. S12.** Phylorate plot showing net diversification rate on a scale from low to high values (blue to red) along each branch of the *Isodon* phylogeny. Red circle represents major rate shift.

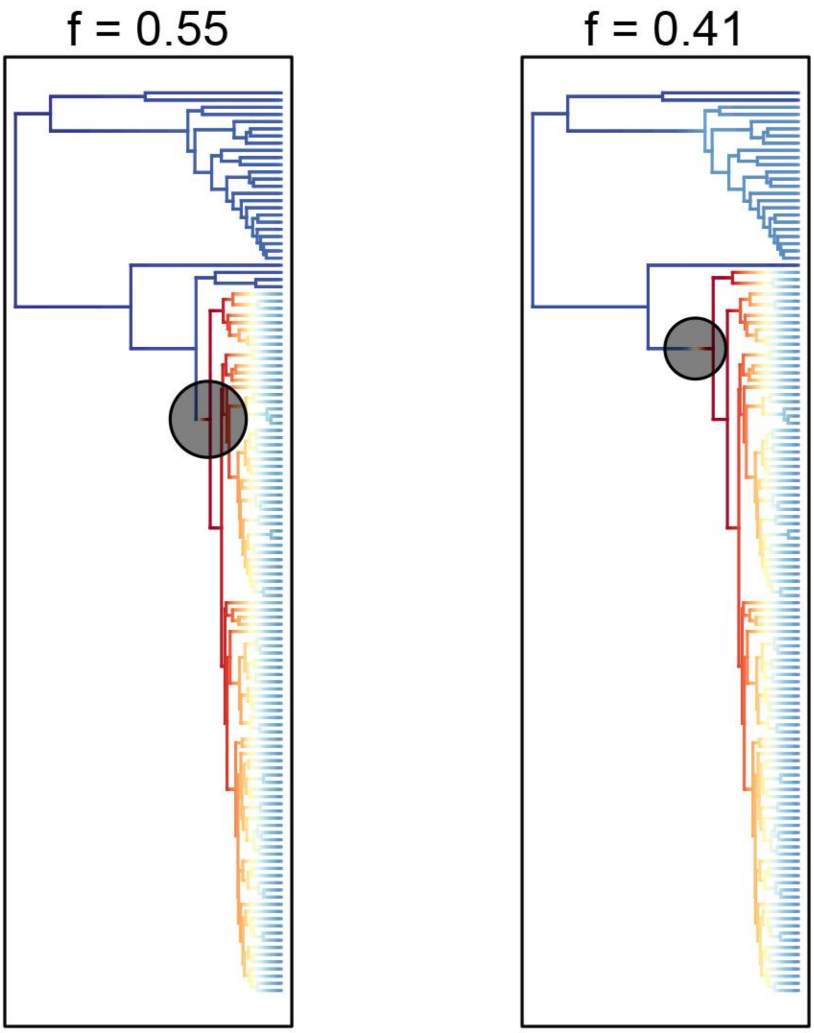


**Fig. S13.** The 95% credible sets of rate shift configurations from BAMM analyses (prior = 1). Significant rate shifts are marked by a circle along the branch at which the shift happened.

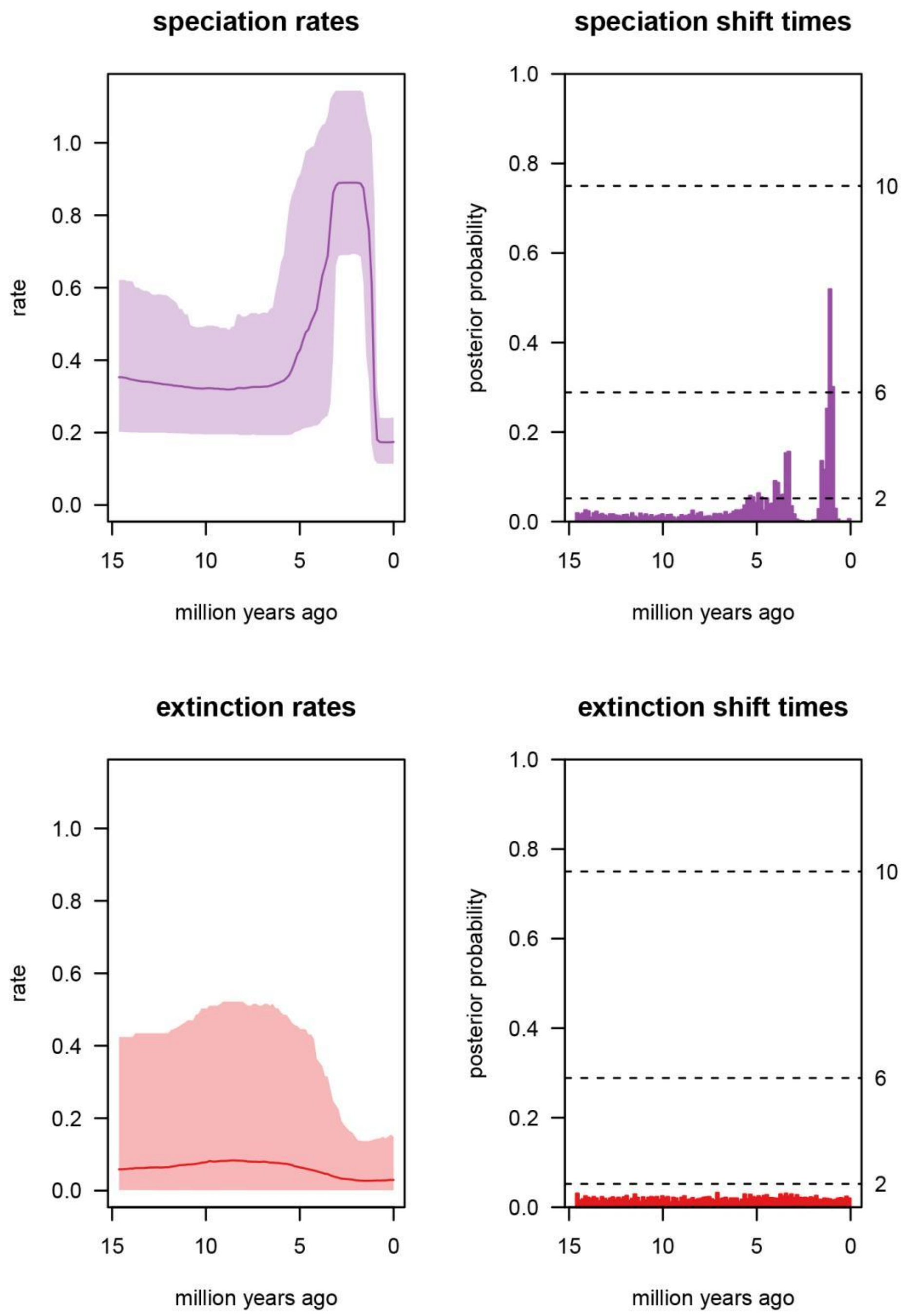


**Fig. S14.** Estimating rates of (and identifying shifts in) lineage diversification through time from TESS analysis. Left: Plots of the posterior mean and 95% credible interval for the speciation and extinction rate (upper and lower panels, respectively). Right: Identifying temporal shifts in the speciation and extinction rate (upper and lower panels, respectively) using Bayes factors (BF) estimated by rjMCMC. Each bar indicates the posterior probability of at least one rate shift within that interval. Bars that exceed the specified significance threshold (2ln BF > 6) indicate significant rate shifts.

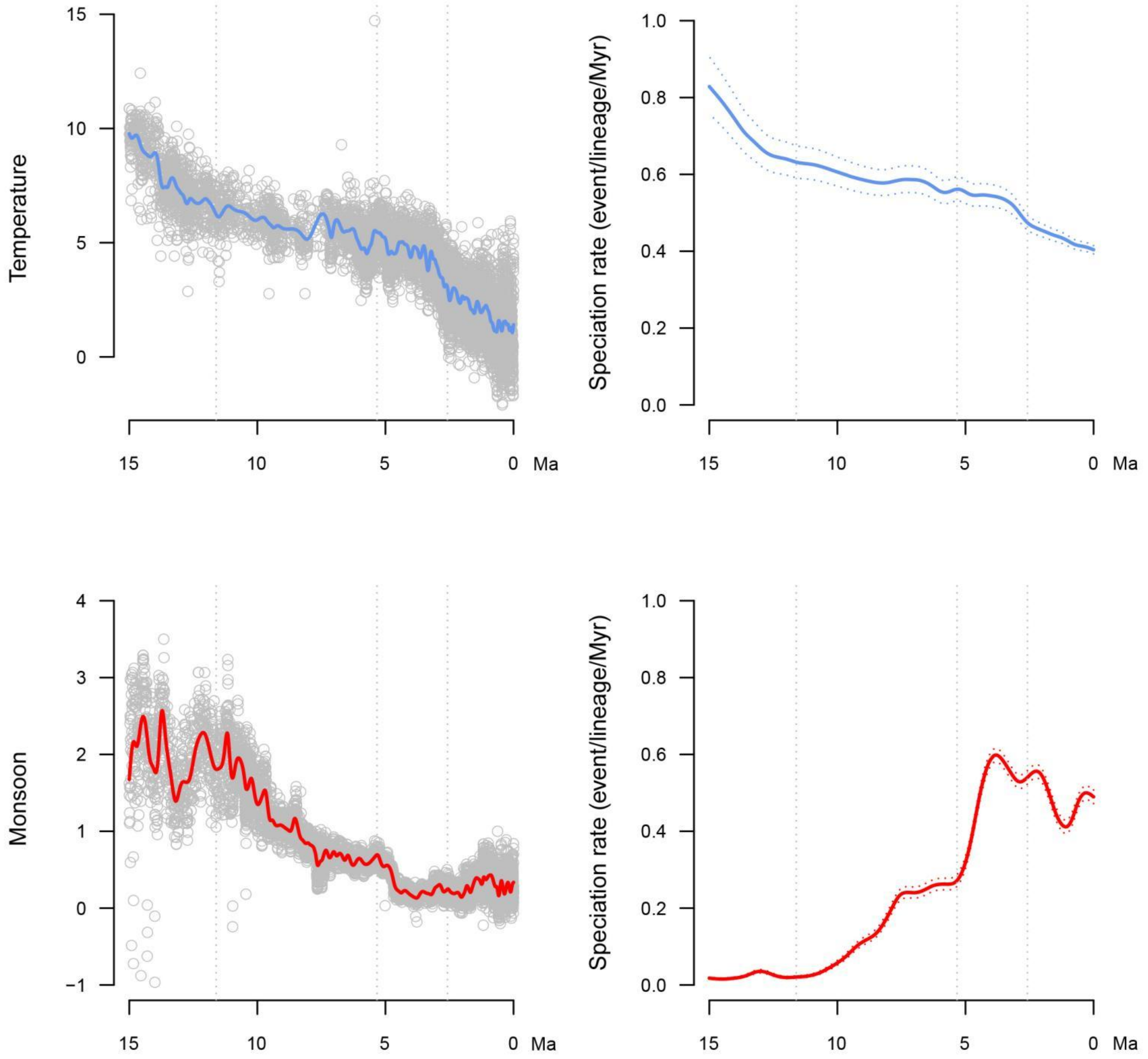


**Fig. S15.** Paleoenvironment-dependent diversification processes in *Isodon*. Left: Past fluctuations of temperature and East Asian monsoon since the middle Miocene (upper and lower panels, respectively). Right: Speciation rates through time for *Isodon* obtained from the relationship between speciation rate and paleotemperature and East Asian monsoon (upper and lower panels, respectively).

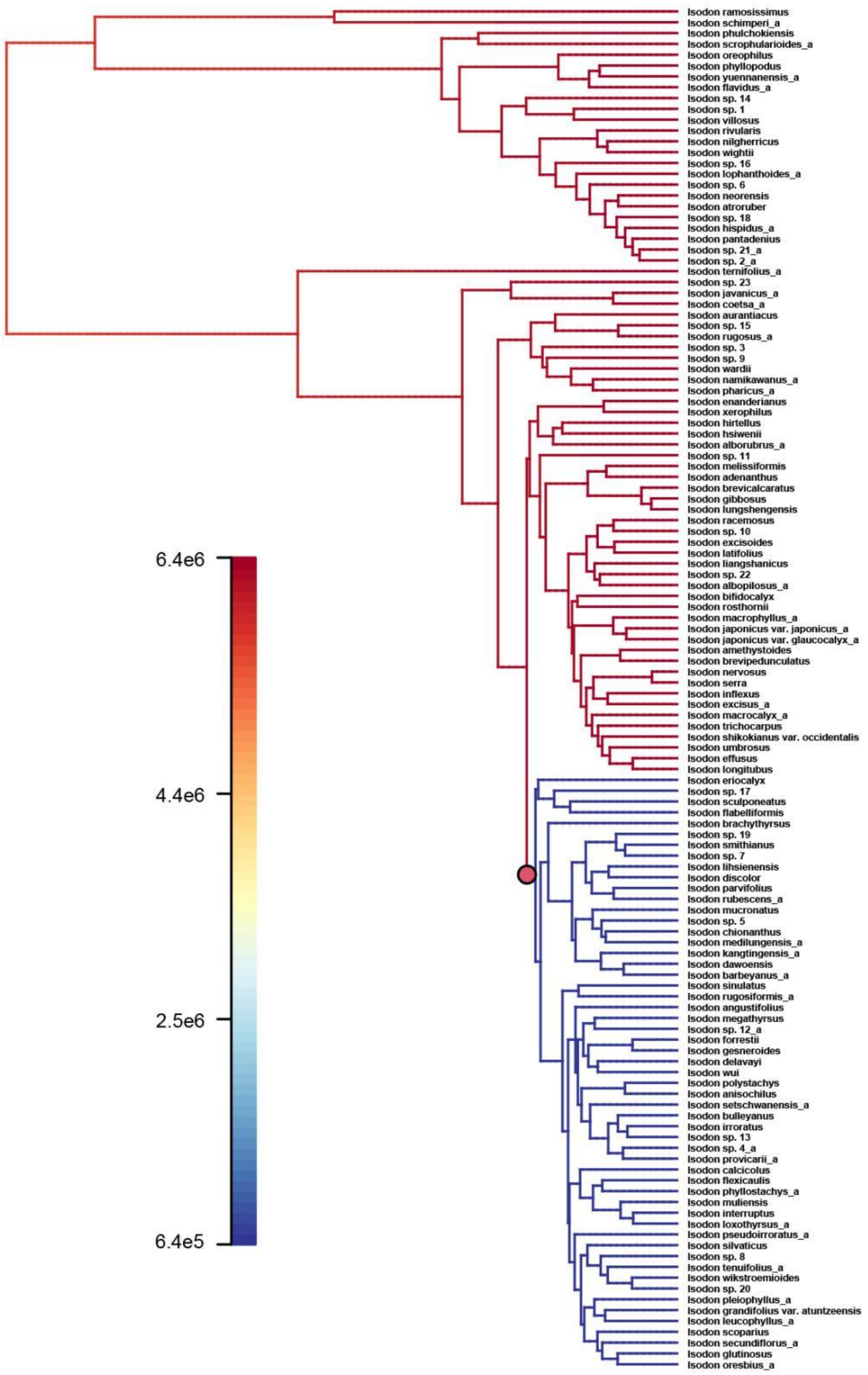


**Fig. S16.** Phylorate plot showing niche rate on a scale from high (red, humid) to low values (blue, arid) along each branch of the *Isodon* phylogeny. Red circle represents major niche shift.

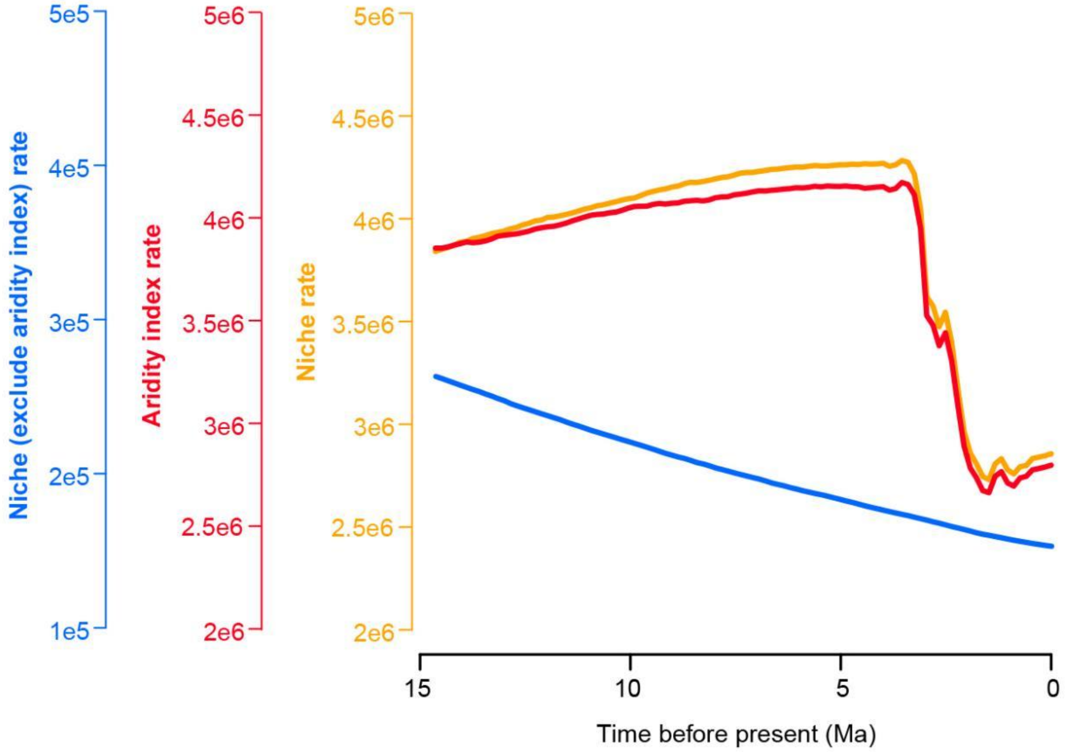


**Fig. S17.** Rates for niche (orange), niche excluding the aridity index (skyblue), and aridity index (red) estimated by BAMM.

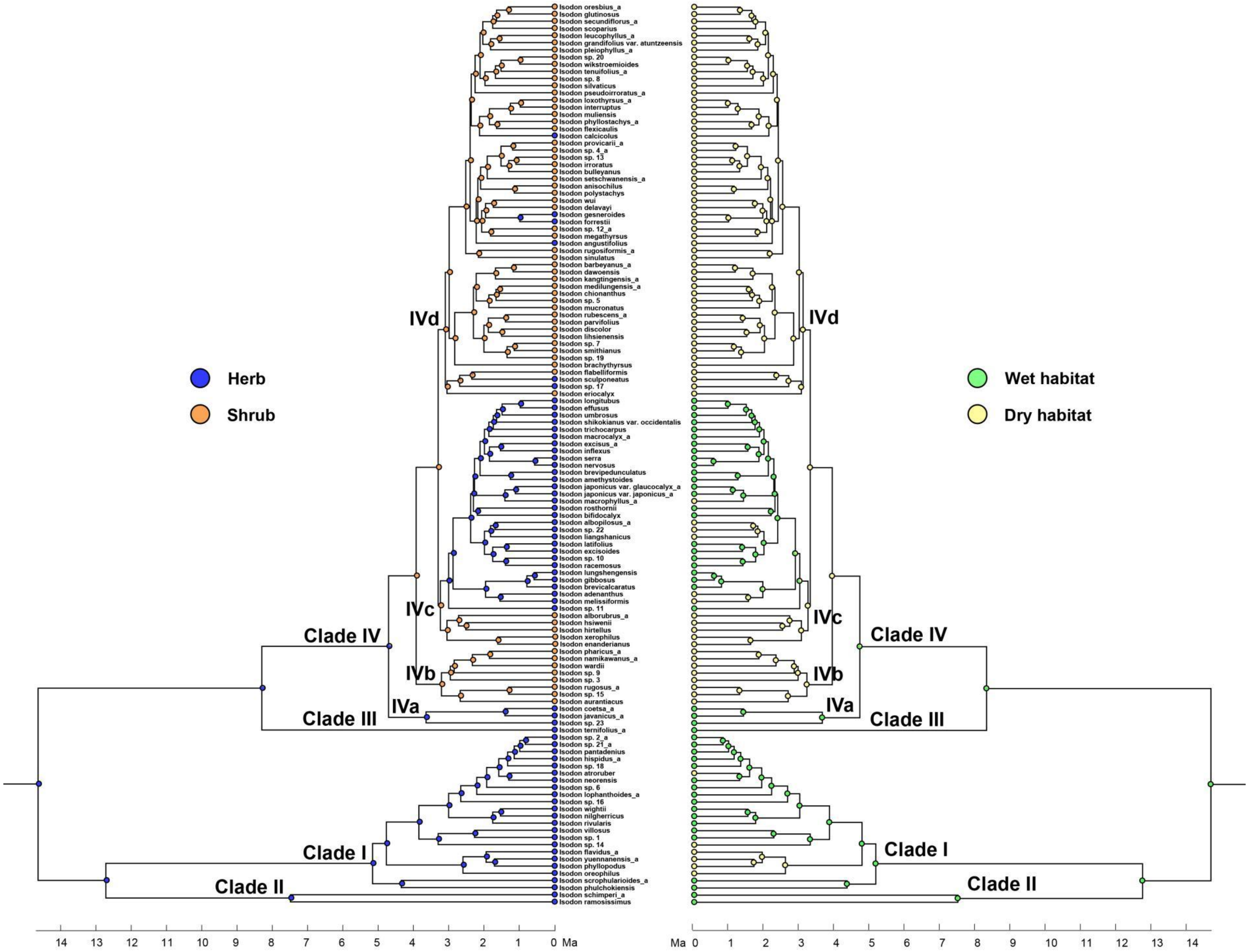


**Fig. S18.** Ancestral state reconstruction of growth form (left) and habitat types (right).

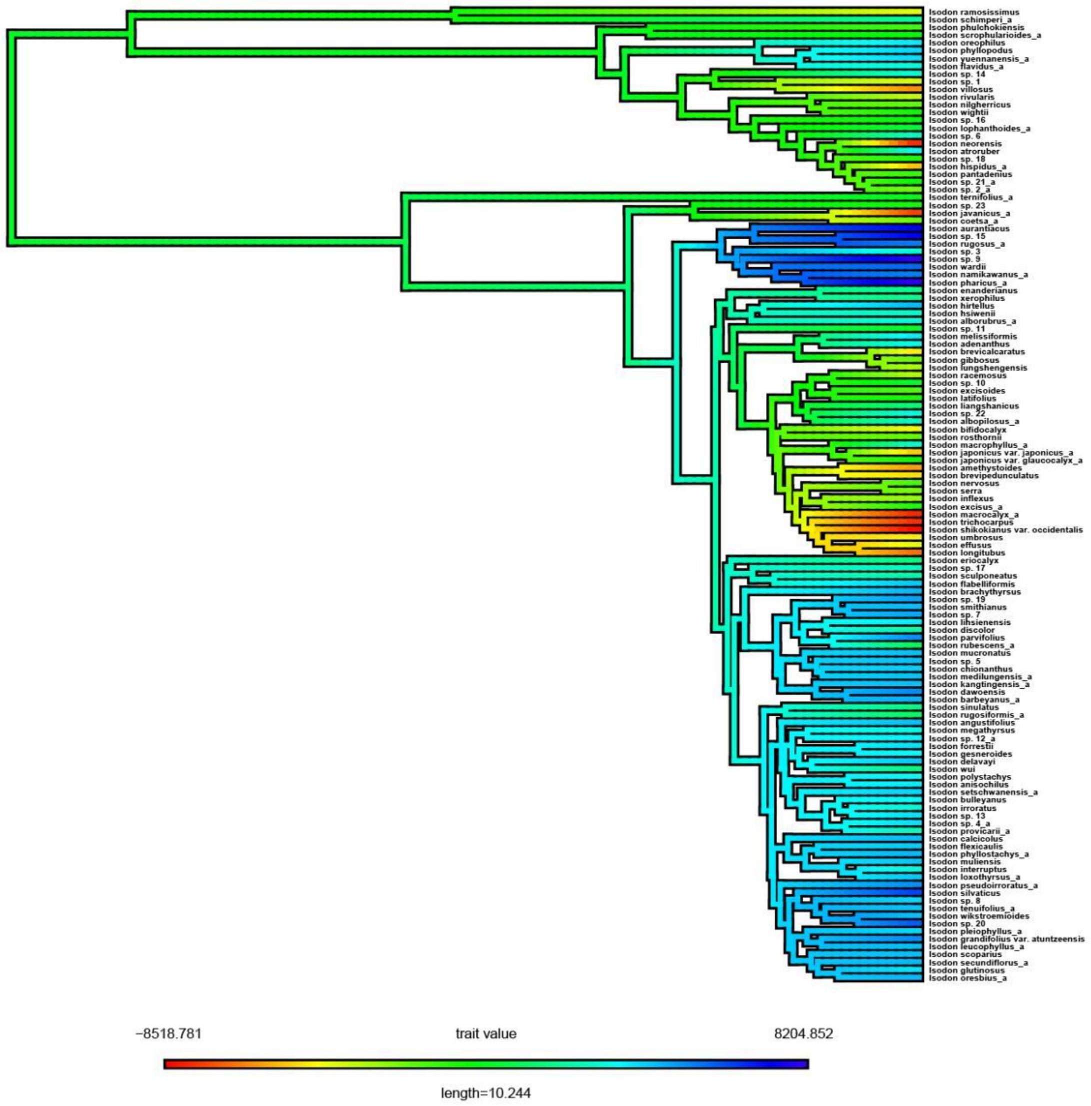


**Fig. S19.** Ancestral reconstruction across *Isodon* for PC1 (Principle Component 1) of the niche dataset. Branches are colored in a rainbow scale from low ordinated values (red and yellow; humid habitats) to high ordinated values (green and blue; arid habitats).

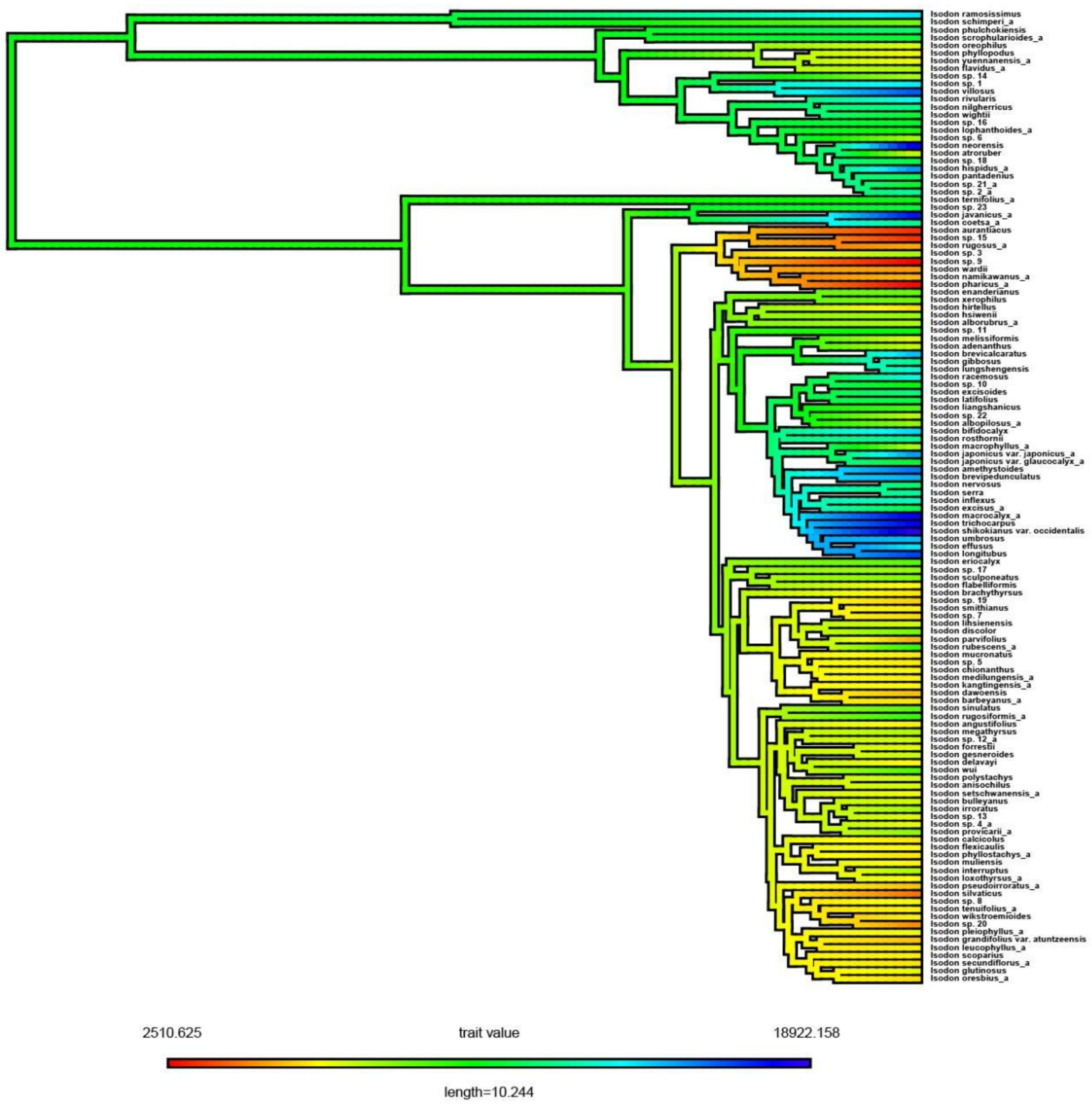


**Fig. S20.** Ancestral reconstruction across *Isodon* for PC1 (Principle Component 1) of the aridity index dataset. Branches are colored in a rainbow scale from low ordinated values (red and yellow; arid habitats) to high ordinated values (green and blue; humid habitats).

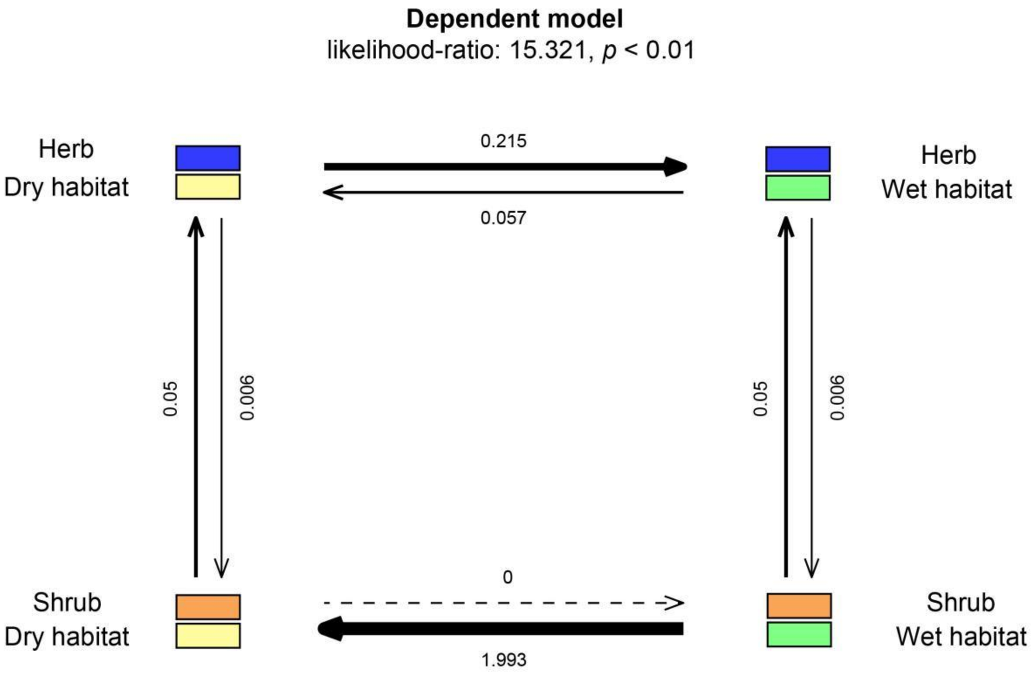


**Fig. S21.** Correlated evolution test of two binary trait, i.e., growth form (herb vs. shrub) and habitat type (wet vs. dry), given four different combination of trait states. Arrows represent the direction of transition and values on arrows indicate transition rates.

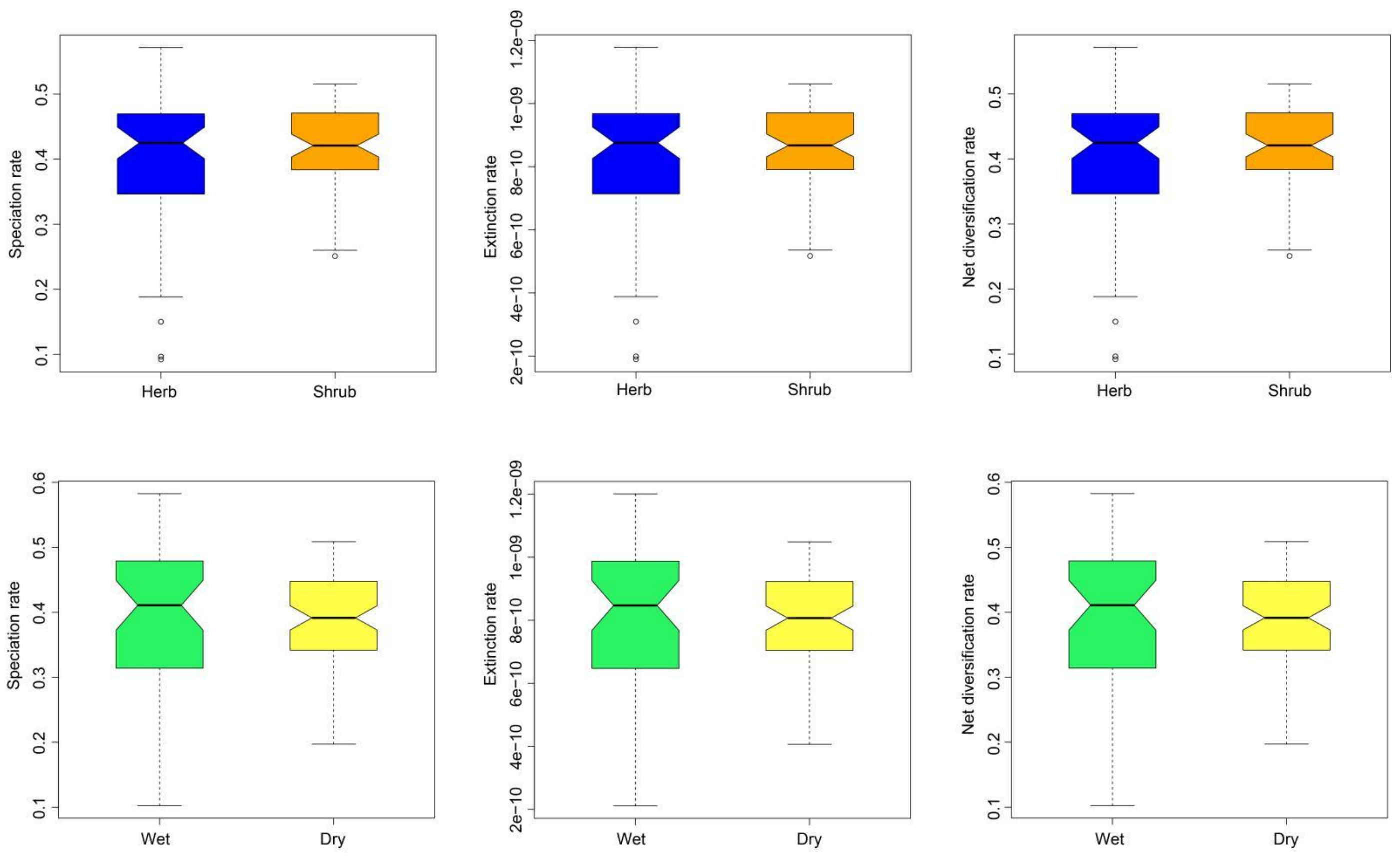


**Fig. S22.** Diversification rate variations of different states for growth form and habitat estimated across tips and nodes after averaging 24 models in HiSSE analyses. The horizontal bars indicate mean values.

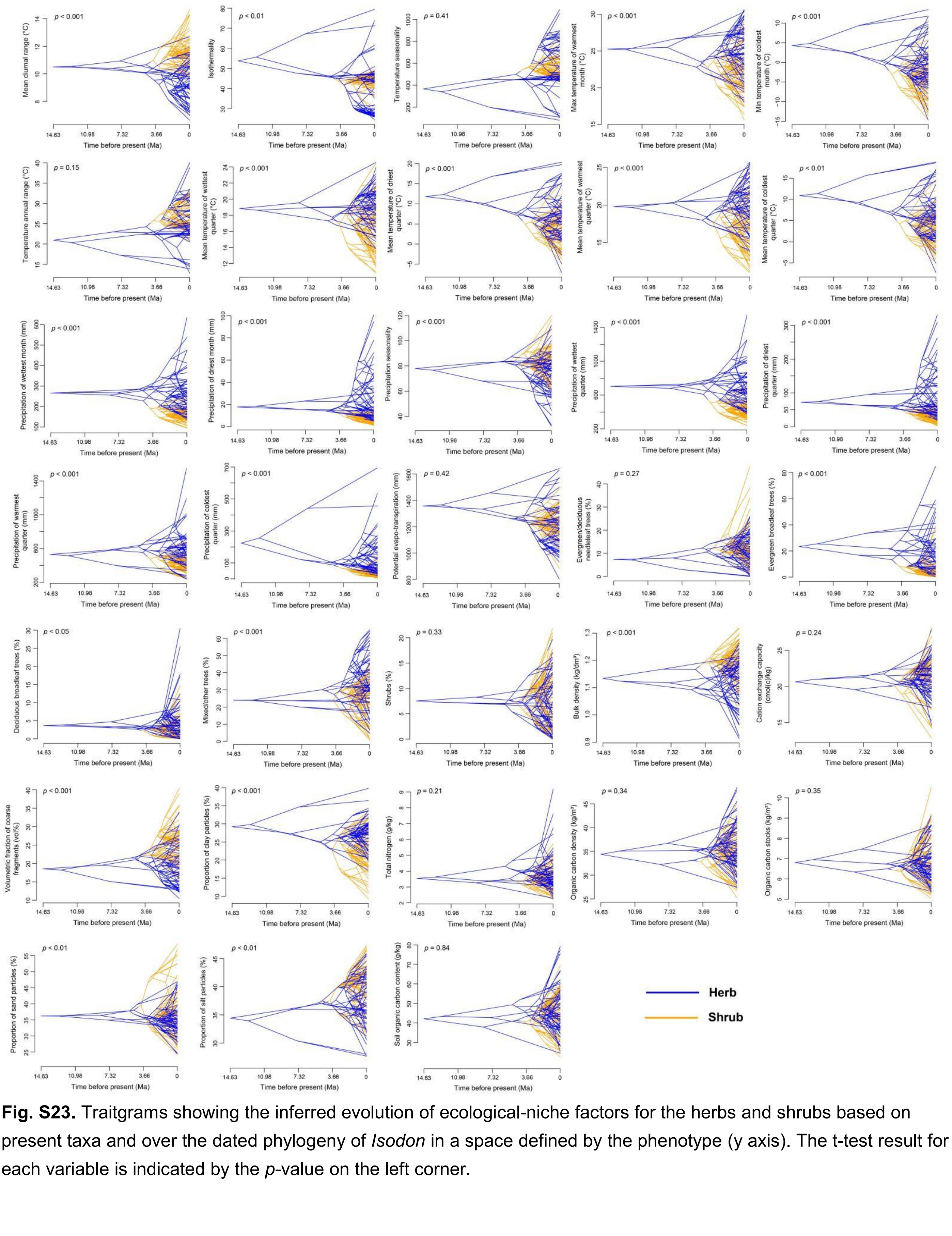


**Fig. S23.** Traitgrams showing the inferred evolution of ecological-niche factors for the herbs and shrubs based on present taxa and over the dated phylogeny of *Isodon* in a space defined by the phenotype (y axis). The t-test result for each variable is indicated by the *p*-value on the left corner.

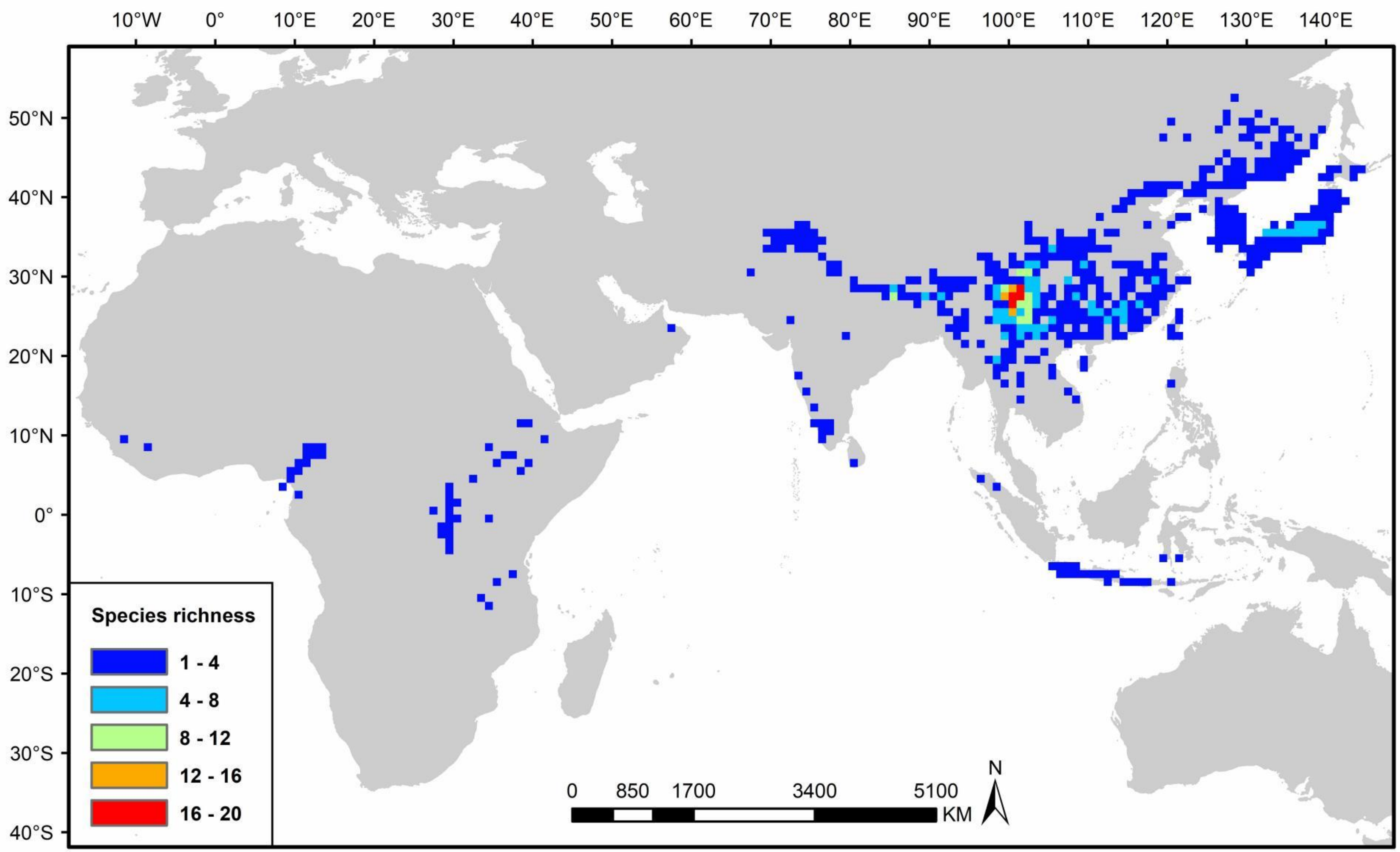


**Fig. S24.** The global patterns of species diversity richness of *Isodon* plotted in grid cells of 100 km × 100 km.

**Table S1.** Biogeographic models tested in this study with estimated parameters from BioGeoBEARS analyses.

| Model | Log-likelihood | No. of parameters | d | e | AICc | AICc_wt |
|---|---|---|---|---|---|---|
| DEC | -144.5 | 2 | 0.026 | 1.00e-12 | 293.2 | 1.00 |
| DIVALIKE | -154.5 | 2 | 0.023 | 2.00e-09 | 313 | 4.9e-05 |
| BAYAREALIKE | -189.1 | 2 | 0.015 | 0.088 | 382.3 | 4.5e-20 |

**Table S2.** Stochastic mapping statistics and data from BioGeoBears results.

Summary of dispersal events counts (mean)

| | A | B | C | D | E | F |
|---|---|---|---|---|---|---|
| A | 0 | 2.1 | 1.44 | 2.28 | 0 | 0.04 |
| B | 1.28 | 0 | 8.18 | 1.5 | 0 | 0.06 |
| C | 0.64 | 2.92 | 0 | 1.4 | 3 | 0.06 |
| D | 2.54 | 1.36 | 2.36 | 0 | 0 | 0.04 |
| E | 0 | 0 | 0 | 0 | 0 | 0 |
| F | 0.02 | 0.06 | 0.06 | 0.02 | 0 | 0 |

Summary counts of 50 BSMs

| | founder | a | d | e | subset | vicariance | sympatry | all_clado | ALL_disp | ana_disp | all_ana | total_events |
|---|---|---|---|---|---|---|---|---|---|---|---|---|
| means | 0 | 0 | 31.36 | 0 | 12.1 (8%) | 11.46 (7%) | 101.4 (65%) | 125 (80%) | 31.36 (20%) | 31.36 (20%) | 31.36 (20%) | 156.4 |
| stdevs | 0 | 0 | 1.14 | 0 | 2.73 | 1.03 | 2.17 | 0 | 1.14 | 1.14 | 1.14 | 1.14 |
| sums | 0 | 0 | 1568 | 0 | 605 | 573 | 5072 | 6250 | 1568 | 1568 | 1568 | 7818 |

A, the Himalaya, together with part of the Hajar Mountains in the Arabian Peninsula; B, the Hengduan Mountains and Yunnan Plateau; C, subtropical China, part of northern China and the Russian Far East, and the Korean Peninsula; D, South and Southeast Asia; E, the Japanese Archipelago; F, Africa. ALL_disp, all dispersal (mean of all observed anagenetic ‘a’, ‘d’ dispersals, plus cladogenetic founder/jump dispersal); ana_disp, Anagenetic dispersal (mean of all observed anagenetic ‘a’ or ‘d’ dispersals); all_ana, all Anagenetic (mean of all observed ‘a’, ‘d’, and ‘e’); all_clado, all Cladogenetic (mean of all sympatry, plus founder, and vicariance).

**Table S3.** Results of RPANDA diversification analyses for *Isodon*. The best-fit model are displayed in bold. Values represent the mean of each parameter as estimated from 200 randomly selected trees of the posterior distribution of the BEAST analyses. Abbreviation: B = birth or speciation; D = death or extinction; NP = number of parameters in the model; AICc = the corrected Akaike information criterion; ΔAICc = the difference in AICc between the model with the lowest AICc and the others; $\lambda_0$ = speciation rate at present; α = shape of speciation rate; $\mu_0$ = extinction rate at present; β = shape of extinction rate.

| Model type | Model description | Rate variation | NP | AICc | ΔAICc | $\lambda_0$ | α | $\mu_0$ | β |
|---|---|---|---|---|---|---|---|---|---|
| Constant rate | Yule model | Constant | 1 | 467.059 | 13.308 | 0.4581 | — | — | — |
| | Constant BD | Exponential | 2 | 469.124 | 15.373 | 0.4581 | — | 0.0000 | — |
| Time dependence | B variable no D | Exponential | 2 | 468.392 | 14.641 | 0.4502 | 0.0111 | — | — |
| | B variable D constant | Exponential | 3 | 470.492 | 16.741 | 0.4502 | 0.0112 | 0.0000 | — |
| | B constant D variable | Exponential | 3 | 471.025 | 17.274 | 0.4589 | — | 0.0000 | 0.0470 |
| | B variable D variable | Exponential | 4 | 468.894 | 15.143 | 0.4271 | 0.0632 | 0.0158 | 0.1691 |
| Temperature dependence | B variable no D | Exponential | 2 | 466.526 | 12.775 | 0.3669 | 0.0831 | — | — |
| | B variable D constant | Exponential | 3 | 468.626 | 14.875 | 0.3674 | 0.0826 | 0.0000 | — |
| | B constant D variable | Exponential | 3 | 471.431 | 17.680 | 0.4566 | — | 0.0000 | 0.0284 |
| | B variable D variable | Exponential | 4 | 470.650 | 16.899 | 0.3653 | 0.0853 | 0.0001 | 0.0611 |
| **Monsoon dependence** | **B variable no D** | **Exponential** | **2** | **453.751** | **0.000** | **0.8155** | **-1.8242** | **—** | **—** |
| | B variable D constant | Exponential | 3 | 455.854 | 2.103 | 0.8098 | -1.8017 | 0.0000 | — |
| | B constant D variable | Exponential | 3 | 471.431 | 17.680 | 0.4565 | — | 0.0000 | 0.0300 |
| | B variable D variable | Exponential | 4 | 457.999 | 4.248 | 0.8045 | -1.7798 | 0.0000 | 0.1543 |

**Table S4.** Niche PCA loadings based on all 39 environmental factors.

| Variables | Abbreviation | PC1 loading |
|---|---|---|
| Aridity index | AI | -0.999749081 |
| Annual precipitation | BIO12 | -0.941287418 |
| Precipitation of wettest quarter | BIO16 | -0.769018383 |
| Precipitation of wettest month | BIO13 | -0.726320761 |
| Precipitation of driest quarter | BIO17 | -0.702828732 |
| Precipitation of driest month | BIO14 | -0.683522518 |
| Precipitation of warmest quarter | BIO18 | -0.631022011 |
| Precipitation of coldest quarter | BIO19 | -0.549433851 |
| Total nitrogen (N) | nitrogen | -0.365746813 |
| Minimum temperature of coldest month | BIO6 | -0.364746741 |
| Organic carbon density | ocd | -0.340291566 |
| Mean temperature of driest quarter | BIO9 | -0.328463086 |
| Soil organic carbon content in the fine earth fraction | soc | -0.321479126 |
| Mean temperature of warmest quarter | BIO10 | -0.298439858 |
| Annual mean temperature | BIO1 | -0.288834672 |
| Evergreen broadleaf trees | consensus2 | -0.286252189 |
| Proportion of clay particles (< 0.002 mm) in the fine earth fraction | clay | -0.269434673 |
| Mean temperature of coldest quarter | BIO11 | -0.268688179 |
| Mixed/other trees | consensus4 | -0.246689478 |
| Maximum temperature of warmest month | BIO5 | -0.232940118 |
| Mean temperature of wettest quarter | BIO8 | -0.209416691 |
| Organic carbon stocks | ocs | -0.188162298 |
| Evergreen/deciduous needleleaf trees | consensus1 | -0.129420616 |
| Cation exchange capacity of the soil | cec | -0.114744125 |
| Proportion of sand particles (> 0.05 mm) in the fine earth fraction | sand | 0.032479868 |
| Deciduous broadleaf trees | consensus3 | 0.041786994 |
| Temperature seasonality (standard deviation ×100) | BIO4 | 0.070227829 |
| Shrubs | consensus5 | 0.160708721 |
| Isothermality (BIO2/BIO7) (×100) | BIO3 | 0.168580716 |
| Potential evapo-transpiration | PET | 0.211929258 |
| Proportion of silt particles (≥ 0.002 mm and ≤ 0.05 mm) in the fine earth fraction | silt | 0.263221916 |
| Temperature annual range (BIO5-BIO6) | BIO7 | 0.304622744 |
| Herbaceous vegetation | consensus6 | 0.349670848 |
| Bulk density of the fine earth fraction | bdod | 0.374704741 |
| Volumetric fraction of coarse fragments (> 2 mm) | cfvo | 0.379384472 |
| Precipitation seasonality (coefficient of variation) | BIO15 | 0.410900779 |
| Elevation | ELEV | 0.419122347 |
| Mean diurnal range (mean of monthly (max temp - min temp)) | BIO2 | 0.60458046 |
| Soil pH | phh2o | 0.624975585 |

**Table S5.** Fit and parameter estimation of four models in the correlated evolution test for two binary traits, i.e., growth form (herb vs. shrub) and habitat type (wet vs. dry). The best-fit model (lowest AIC_weight) is marked in bold.

| Model | Log-likelihood | AIC | AIC_weight |
|---|---|---|---|
| Independent | -64.68 | 137.37 | 0.00 |
| Dependent (growth form and habitat) | -57.34 | 130.68 | 0.07 |
| Dependent (growth form) | -58.55 | 129.10 | 0.16 |
| **Dependent (habitat)** | **-57.02** | **126.05** | **0.76** |

**Table S6.** Fit and parameter estimation of 24 speciation and extinction models with hidden traits (HiSSE) fitted to growth form and habitat. τ = turnover rate; ε = extinction fraction; *q* = transition rates. All states = 0A, 0B, 0C, 0D, 1A, 1B, 1C, 1D (numbers represent "known" states and letters represent "hidden" states); state 0 = herb or wet habitat; state 1 = shrub or dry habitat. The best-fit model (lowest AICc) is marked in bold.

| Model | growth form | | | | Habitat | | | |
|---|---|---|---|---|---|---|---|---|
| | Log-likelihood | AIC | AICc | ΔAICc | Log-likelihood | AIC | AICc | ΔAICc |
| BiSSE1<br>τ = 0 ≠ 1<br>ε = 0 ≠ 1<br>$q_{observed}$ = 0→1 = 1 →0 | -262.42 | 534.84 | 535.34 | 26.61 | -269.72 | 549.45 | 549.95 | 34.43 |
| BiSSE2<br>τ = 0 ≠ 1<br>ε = 0 = 1<br>$q_{observed}$ = 0→1 = 1 →0 | -262.42 | 532.84 | 533.17 | 24.44 | -269.76 | 547.51 | 547.84 | 32.32 |
| HiSSE-CID-2<br>τ = (0A = 1A) ≠ (0B = 1B)<br>ε = (0A = 1A) ≠ (0B = 1B)<br>*q* = all equal | -256.32 | 522.64 | 523.14 | 14.41 | -262.52 | 535.04 | 535.54 | 20.02 |
| HiSSE-CID-2-constrained<br>τ = (0A = 1A) ≠ (0B = 1B)<br>ε = all equal<br>*q* = all equal | -256.32 | 520.64 | 520.97 | 12.24 | -262.52 | 533.04 | 533.37 | 17.85 |
| HiSSE-CID-4<br>τ = (0A = 1A) ≠ (0B = 1B) ≠ (0C = 1C) ≠ (0D = 1D)<br>ε = (0A = 1A) ≠ (0B = 1B) ≠ (0C = 1C) ≠ (0D = 1D)<br>*q* = all equal | -248.01 | 514.02 | 515.57 | 6.84 | -251.41 | 520.81 | 522.36 | 6.84 |
| **HiSSE-CID-4-constrained<br>τ = (0A = 1A) ≠ (0B = 1B) ≠ (0C = 1C) ≠ (0D = 1D)<br>ε = all equal<br>*q* = all equal** | **-248.01** | **508.02** | **508.73** | **0** | **-251.41** | **514.81** | **515.52** | **0** |
| HiSSE7<br>τ = 0A ≠1A ≠ 0B ≠1B<br>ε = 0A ≠1A ≠ 0B ≠1B<br>*q* = all equal | -256.03 | 530.05 | 531.60 | 22.87 | -261.94 | 541.89 | 543.44 | 27.92 |
| HiSSE8<br>τ = 0A ≠ 1A ≠ 0B ≠ 1B<br>ε = all equal | -256.03 | 524.05 | 524.76 | 16.03 | -261.94 | 535.89 | 536.59 | 21.07 |

| | | | | | | | | |
|---|---|---|---|---|---|---|---|---|
| *q* = all equal | | | | | | | | |
| HiSSE9<br>τ = (0A = 1A = 0B) ≠ 1B<br>ε = (0A = 1A = 0B) ≠ 1B<br>*q* = all equal | -262.80 | 535.60 | 536.10 | 27.37 | -269.86 | 549.72 | 550.22 | 34.70 |
| HiSSE10<br>τ = (0A = 1A = 0B) ≠ 1B<br>ε = all equal<br>*q* = all equal | -262.80 | 533.60 | 533.93 | 25.20 | -269.86 | 547.72 | 548.05 | 32.53 |
| HiSSE11<br>τ = 0A ≠ 1A ≠ 0B ≠ 1B<br>ε = 0A ≠ 1A ≠ 0B ≠ 1B<br>*q* = all equal<br>*q0A→ q1B, q1A→ q0B, q0B→ q1A, q1B→ q0A = 0* | -258.00 | 534.00 | 535.56 | 26.83 | -261.21 | 540.42 | 541.97 | 26.45 |
| HiSSE12<br>τ = 0A ≠ 1A ≠ 0B ≠ 1B<br>ε = all equal<br>*q* = all equal<br>*q0A→ q1B, q1A→ q0B, q0B→ q1A, q1B→ q0A = 0* | -255.90 | 523.80 | 524.51 | 15.78 | -261.21 | 534.42 | 535.12 | 19.60 |
| HiSSE13<br>τ = (0A = 1A = 0B) ≠ 1B<br>ε = (0A = 1A = 0B) ≠ 1B<br>*q* = all equal<br>*q0A→ q1B, q1A→ q0B, q0B→ q1A, q1B→ q0A = 0* | -262.36 | 534.73 | 535.23 | 26.50 | -269.57 | 549.14 | 549.64 | 34.12 |
| HiSSE14<br>τ = (0A = 1A = 0B) ≠ 1B<br>ε = all equal<br>*q* = all equal<br>*q0A→ q1B, q1A→ q0B, q0B→ q1A, q1B→ q0A = 0* | -262.36 | 532.73 | 533.06 | 24.33 | -269.57 | 547.14 | 547.47 | 31.95 |
| HiSSE15<br>τ = (0A = 0B) ≠ 1A ≠ 1B<br>ε = (0A = 0B) ≠ 1A ≠ 1B<br>*q* = all equal | -262.15 | 538.30 | 539.25 | 30.52 | -269.34 | 552.68 | 553.63 | 38.11 |
| HiSSE16<br>τ = (0A = 0B) ≠ 1A ≠ 1B<br>ε = all equal<br>*q* = all equal | -262.15 | 534.30 | 534.80 | 26.07 | -269.35 | 548.70 | 549.20 | 33.68 |

| | | | | | | | | |
|---|---|---|---|---|---|---|---|---|
| HiSSE17<br>τ = (0A = 0B) ≠1A ≠1B<br>ε = (0A = 0B) ≠1A ≠1B<br>*q* = all equal<br>*q0A→ q1B, q1A→ q0B, q0B→ q1A, q1B→ q0A = 0* | -261.41 | 536.82 | 537.77 | 29.04 | -268.35 | 550.71 | 551.66 | 36.14 |
| HiSSE18<br>τ = (0A = 0B) ≠ 1A ≠ 1B<br>ε = all equal<br>*q* = all equal<br>*q0A→ q1B, q1A→ q0B, q0B→ q1A, q1B→ q0A = 0* | -261.41 | 532.82 | 533.32 | 24.59 | -268.34 | 546.67 | 547.17 | 31.65 |
| HiSSE19<br>τ = (0A = 1A ) ≠ 0B ≠ 1B<br>ε = (0A = 1A ) ≠ 0B ≠ 1B<br>*q* = all equal | -256.06 | 526.12 | 527.07 | 18.34 | -262.34 | 538.68 | 539.63 | 24.11 |
| HiSSE20<br>τ = (0A = 1A) ≠ 0B ≠ 1B<br>ε = all equal<br>*q* = all equal | -256.06 | 522.12 | 522.62 | 13.89 | -262.34 | 534.68 | 535.18 | 19.66 |
| HiSSE21<br>τ = (0A = 1A) ≠ 0B ≠1B<br>ε = (0A = 1A) ≠ 0B ≠1B<br>*q* = all equal<br>*q0A→ q1B, q1A→ q0B, q0B→ q1A, q1B→ q0A = 0* | -256.05 | 526.10 | 527.05 | 18.32 | -261.38 | 536.75 | 537.70 | 22.18 |
| HiSSE22<br>τ = (0A = 1A ) ≠ 0B ≠ 1B<br>ε = all equal<br>*q* = all equal<br>*q0A→ q1B, q1A→ q0B, q0B→ q1A, q1B→ q0A = 0* | -256.05 | 522.10 | 522.60 | 13.87 | -261.38 | 532.75 | 533.25 | 17.73 |
| HiSSE23<br>τ = 0A ≠ 1A ≠ 1B; 0B = 0<br>ε = 0A ≠ 1A ≠ 1B; 0B = 0<br>*q* = all equal | -261.63 | 537.26 | 538.21 | 29.49 | -264.35 | 542.69 | 543.64 | 28.12 |
| HiSSE24<br>τ = 0A ≠ 1A ≠ 1B; 0B = 0<br>ε = 0A ≠ 1A ≠ 1B; 0B = 0<br>*q* = all equal<br>*q0A→ q1B, q1A→ q0B, q0B→ q1A, q1B→ q0A = 0* | -259.08 | 532.15 | 533.10 | 24.37 | -267.29 | 548.59 | 549.54 | 34.02 |

8

**Table S7.** Fit and parameter estimation of Multistate Speciation and Extinction with hidden traits (MuHiSSE) models fitted for *Isodon* and two traits (growth form and habitat type affiliation). τ = turnover rate; ε = extinction fraction; *q* = transition rates. The best-fit model (lowest AICc) is marked in bold.

| Model | Log-likelihood | AIC | AICc | ΔAICc |
|---|---|---|---|---|
| Null model<br>τ = 00 = 01 = 11<br>ε = all equal<br>$q_{observed}$ = all different<br>*q00→ q11, q11→ q00 = 0* | -290.65 | 593.30 | 594.00 | 62.17 |
| MuSSE1<br>τ = 00 ≠ 01 ≠ 11<br>ε = all equal<br>$q_{observed}$ = all different<br>*q00→ q11, q11→ q00 = 0* | -285.13 | 594.26 | 597.02 | 65.19 |
| MuSSE2<br>τ = 00 ≠ 01 ≠ 11<br>ε = 00 ≠ 01 ≠ 11<br>$q_{observed}$ = all different<br>*q00→ q11, q11→ q00 = 0* | -286.76 | 601.52 | 605.31 | 73.48 |
| Full MuHiSSE1<br>τ = 00A ≠ 01A ≠ 11A ≠ 00B ≠ 01B ≠ 11B<br>ε = all equal<br>$q_{observed}$ = all different<br>$q_{hidden}$ = all equal<br>*q00→ q11, q11→ q00 = 0* | -267.03 | 566.06 | 571.05 | 39.22 |
| Full MuHiSSE2<br>τ = 00A ≠ 01A ≠ 11A ≠ 00B ≠ 01B ≠ 11B<br>ε = 00A ≠ 01A ≠ 11A ≠ 00B ≠ 01B ≠ 11B<br>$q_{observed}$ = all different<br>$q_{hidden}$ = all equal<br>*q00→ q11, q11→ q00 = 0* | -264.44 | 570.89 | 579.77 | 47.94 |
| MuHiSSE-CID-2<br>τ = (00A = 01A = 11A) ≠ (00B = 01B = 11B)<br>ε = all equal<br>$q_{observed}$ = all different<br>$q_{hidden}$ = all equal<br>*q00→ q11, q11→ q00 = 0* | -268.13 | 552.25 | 553.49 | 21.66 |
| MuHiSSE-CID-3<br>τ = (00A = 01A = 11A) ≠ (00B = 01B = 11B) ≠ (00C = 01C = 11C)<br>ε = all equal<br>$q_{observed}$ = all different<br>$q_{hidden}$ = all equal<br>*q00→ q11, q11→ q00 = 0* | -259.47 | 536.94 | 538.49 | 6.66 |
| **MuHiSSE-CID-4**<br>**τ = (00A = 01A = 11A) ≠ (00B = 01B = 11B) ≠** | **-254.96** | **529.91** | **531.83** | **0** |

**(00C = 01C = 11C) ≠ (00D = 01D = 11D)**
**ε = all equal**
**$q_{observed}$ = all different**
**$q_{hidden}$ = all equal**
***q00→ q11, q11→ q00 = 0***

13

**Table S8.** The 39 environmental variables used for niche and species richness analysis in this study.

| Abbreviation | Variable description | Unit |
|---|---|---|
| bdod | Bulk density of the fine earth fraction | $cg/cm^3$ |
| cec | Cation exchange capacity of the soil | mmol(c)/kg |
| cfvo | Volumetric fraction of coarse fragments (> 2 mm) | $cm^3/dm^3$ (vol‰) |
| clay | Proportion of clay particles (< 0.002 mm) in the fine earth fraction | g/kg |
| nitrogen | Total nitrogen (N) | cg/kg |
| ocd | Organic carbon density | $hg/m^3$ |
| ocs | Organic carbon stocks | t/ha |
| phh2o | Soil pH | pH x 10 |
| sand | Proportion of sand particles (> 0.05 mm) in the fine earth fraction | g/kg |
| silt | Proportion of silt particles (≥ 0.002 mm and ≤ 0.05 mm) in the fine earth fraction | g/kg |
| soc | Soil organic carbon content in the fine earth fraction | dg/kg |
| consensus1 | Evergreen/deciduous needleleaf trees | % |
| consensus2 | Evergreen broadleaf trees | % |
| consensus3 | Deciduous broadleaf trees | % |
| consensus4 | Mixed/other trees | % |
| consensus5 | Shrubs | % |
| consensus6 | Herbaceous vegetation | % |
| PET | Potential evapo-transpiration | mm |
| AI | Aridity index | |
| BIO1 | Annual mean temperature | °C |
| BIO2 | Mean diurnal range (mean of monthly (max temp - min temp)) | °C |
| BIO3 | Isothermality (BIO2/BIO7) (×100) | |
| BIO4 | Temperature seasonality (standard deviation ×100) | |
| BIO5 | Maximum temperature of warmest month | °C |
| BIO6 | Minimum temperature of coldest month | °C |
| BIO7 | Temperature annual range (BIO5-BIO6) | °C |
| BIO8 | Mean temperature of wettest quarter | °C |
| BIO9 | Mean temperature of driest quarter | °C |
| BIO10 | Mean temperature of warmest quarter | °C |
| BIO11 | Mean temperature of coldest quarter | °C |
| BIO12 | Annual precipitation | mm |
| BIO13 | Precipitation of wettest month | mm |
| BIO14 | Precipitation of driest month | mm |
| BIO15 | Precipitation seasonality (coefficient of variation) | |
| BIO16 | Precipitation of wettest quarter | mm |
| BIO17 | Precipitation of driest quarter | mm |
| BIO18 | Precipitation of warmest quarter | mm |
| BIO19 | Precipitation of coldest quarter | mm |
| ELEV | Elevation | m |

**Table S9.** Summary of the results of multi-predictor regression models explaining the species richness of *Isodon* within 100 km × 100 km grid cells. $R^2$, the explained variance of the environmental variables. Significance levels: ***$p$ < 0.001; **$p$ < 0.01; *$p$ < 0.05. Abbreviations and explanations of predictor variables refer to *SI Appendix*, Table S8.

| | Estimate coefficient | Standard error | *t*-value | *p*-value | Significance level |
|---|---|---|---|---|---|
| Intercept | 0.94313 | 0.03632 | 25.967 | < 2e-16 | *** |
| consensus1 | 0.27374 | 0.03886 | 7.044 | 5.44e-12 | *** |
| cfvo | 0.25389 | 0.04125 | 6.156 | 1.42e-09 | *** |
| BIO1 | 0.30073 | 0.04985 | 6.033 | 2.91e-09 | *** |
| ocs | 0.25185 | 0.04134 | 6.092 | 2.06e-09 | *** |
| silt | 0.12994 | 0.04253 | 3.056 | 0.00235 | ** |
| BIO19 | -0.14569 | 0.0446 | -3.266 | 0.00116 | ** |
| consensus3 | -0.08959 | 0.03787 | -2.366 | 0.01834 | * |
| $R^2$ | 0.2426 | | | < 2.2e-16 | *** |
